# Genome-wide functional genomics screens identify therapeutic vulnerabilities for lymphangioleiomyomatosis

**DOI:** 10.1101/683359

**Authors:** Alberto Camacho-Magallanes, Sean P. Delaney, Eric Lian, Ellis Wiljer, Julien Yockell-Lelièvre, Lisa M. Julian, Adam Pietrobon, William Batoff, Arnold S Kristof, William L. Stanford

**Author notes:** **Correspondence:** WLS. Equal contribution (Alberto Camacho-Magallanes, Sean P. Delaney).

## Abstract

Lymphangioleiomyomatosis (LAM) is a rare lung disease marked by cystic destruction caused by invasive LAM cells harboring loss-of-function mutations in *TSC2* that exhibit dysregulated mTORC1 signaling. Rapamycin, the only approved treatment, is not curative. Therapeutic discovery has been limited by inadequate cell and animal models due to the unknown LAM cell of origin. We report a novel *TSC2^—/—^* human pluripotent stem cell (hPSC)-derived neural crest cell (NCC) model, which replicates key LAM features. Leveraging this model, temporal RNA sequencing and genome-wide CRISPR knockout screens identified synthetic lethal genes including *TWIST1* and *SUN4*. Antisense oligonucleotides (ASOs) targeting TWIST1 and SUN4 demonstrated selective TSC2 mutant NCC cytotoxicity and inhibited invasion. Using a “test-withdraw-monitor” clinical trial paradigm, TWIST1 and SUN4 ASOs profoundly suppressed tumor viability in a *Tsc2^—/—^* mouse allograft model and eradicated 40% of tumors, outperforming standard of care rapamycin and highlighting their promise as novel cytoablative LAM therapeutics.

## Introduction

Lymphangioleiomyomatosis (LAM) is a rare cystic lung disease, almost exclusively affecting women, with an estimated prevalence of 25 per million females^1^. LAM occurs sporadically or in association with tuberous sclerosis complex (TSC), a monogenic multisystem tumor-forming disorder^2,3^. LAM is characterized by the presence of neoplastic lesions within the lung interstitium that share molecular and histopathological features with extrapulmonary TSC-associated tumors, including the kidney (renal angiomyolipoma, rAML)^4–6^, uterus (uterine PEComas)^7^, and lymphatics (lymphangioleiomyomas)^8^. LAM nodules consist of non-transformed, slowly proliferating smooth muscle-like cells and abnormally large epithelial cells expressing the smooth muscle marker ACTA2/αSMA and melanocytic markers (PMEL, MLANA)^9^, originating from an unknown cell of origin. LAM clinical features are characterized by progressive dyspnea, chylous pleural effusions, pneumothoraces, and progressive parenchymal lung destruction leading to respiratory failure ^10–13^.

The molecular etiology associated with LAM and related TSC-associated tumors involves loss of function mutations in *TSC2*, or, less commonly, *TSC1*. Together with TBC1D7, TSC1 and TSC2 form the TSC complex, which negatively regulates mechanistic target of rapamycin complex 1 (mTORC1)^14,15^. Loss of TSC function disrupts mTORC1 regulation, resulting in dysregulated mTORC1 signaling^14,16^. These insights led to clinical trials and the eventual approval of rapamycin (clinically known as sirolimus, everolimus) as the only approved treatment for LAM and other TSC-associated manifestations, including rAML^17,18^. Although daily rapamycin treatment reduces the rate of lung function decline in most LAM patients, others experience partial or no response^19,20^. Rapamycin is not curative due to its cytostatic and cytoprotective effects, with disease progression occurring rapidly upon treatment withdrawal^17,18^. Notably, several LAM and TSC tumor features are mTORC1 independent and remain unaffected by rapamycin treatment^21–23^. These limitations highlight the need to identify cytocidal therapeutics that eradicate LAM cells.

Mechanistic characterization of mTORC1-independent phenotypes and therapeutic development have been limited by the lack of cell and animal models that replicate the complexity of LAM. Recent single-cell RNA-sequencing (RNA-seq) studies confirmed findings from histological analyses that LAM cells represent a minority of cells within the LAM nodule^24–27^. In addition to the limited availability of viable primary specimens, the heterogeneous nature of LAM nodules has impeded the establishment of stable, homogenous populations of LAM cells^28^. We previously generated female *TSC2^—/—^* smooth muscle cells (SMCs) to model the SMC component of LAM ^23,29,30^. This non-transformed human cell-based model enabled the identification of histone deacetylase (HDAC) inhibitors as a class of drugs with both anti-invasive activity and *TSC2^—/—^*specific cytotoxicity^23^. While this model recapitulated several LAM-relevant phenotypes, only a small fraction of cells (approximately 1%) expressed the pathognomonic marker PMEL^23’^. These findings highlight the need for models that more fully recapitulate the molecular and phenotypic features of LAM cells, including characteristics shared across pulmonary and extrapulmonary manifestations of TSC-associated lesions.

Neural crest cells (NCCs) offer a compelling framework for modeling LAM-like cells. NCCs are multipotent, highly migratory, and give rise to cell types relevant to LAM pathology, including both smooth muscle and melanocytic lineages^2^. Moreover, modeling rAML using *TSC1^—/—^* and *TSC2^—/—^* kidney organoids identified neural crest-derived glial cells as a progenitor population capable of generating PMEL-positive cells^23^.

We present a human pluripotent stem cell (hPSC)-derived NCC model that demonstrates key histopathological and transcriptional features of LAM cells, providing a relevant platform for therapeutic discovery. To uncover novel therapeutic targets that could selectively eradicate LAM cells, we conducted a genome-wide CRISPR/Cas9-mediated synthetic lethal screen using wild-type (WT) and *TSC2^—/—^* NCCs. This approach identified *TWIST1* and *SUN4* (also known as SPAG4) as negatively selected targets, specifically in *TSC2^—/—^* cultures. Antisense oligonucleotides (ASOs) targeting TWIST1 and SUN4 in our NCC lines and *in vivo* using a TSC allograft model confirmed their selective cytotoxicity, anti-invasiveness, and anti-tumorigenicity towards *TSC2^—/—^*cells, with a proportion of tumors entirely eradicated. Critically, these ASOs induce apoptosis in the presence of rapamycin, suggesting they could be used in combination with rapamycin. Thus, these findings identify TWIST1 and SUN4 as candidate therapeutic targets and support the potential of non–mTORC1–directed strategies that selectively eradicate pathogenic LAM cells and outperform the current standard of care, rapamycin, in preclinical models of LAM.

## Results

### Generation of hPSC library to model TSC

To model LAM and other TSC-related pathologies, CRISPR-based homology-directed repair was used to generate identical mutations in exon 3 of the *TSC2* locus in four genetically distinct wild-type (WT) hPSC lines, composed of two male (H1 & D168) and two female (H7 & H9) lines **(Figure 1A)**. Exon 3 was selected for targeted disruption due to its presence in all known *TSC2* isoforms and its proximity to the N-terminus of its protein product. Additionally, each parental line was modified at the AAVS1 locus with CRISPR to express green (eGFP) or red (mCherry) fluorescent reporters along with Zeocin resistance to aid in live cell assays and future *in vivo* studies. The TSC2 knockout strategy inserted premature stop codons in all three reading frames, as confirmed by PmeI digestion of PCR amplicons spanning the targeted site **(Figure 1B)** and Sanger sequencing.

**Figure 1:**
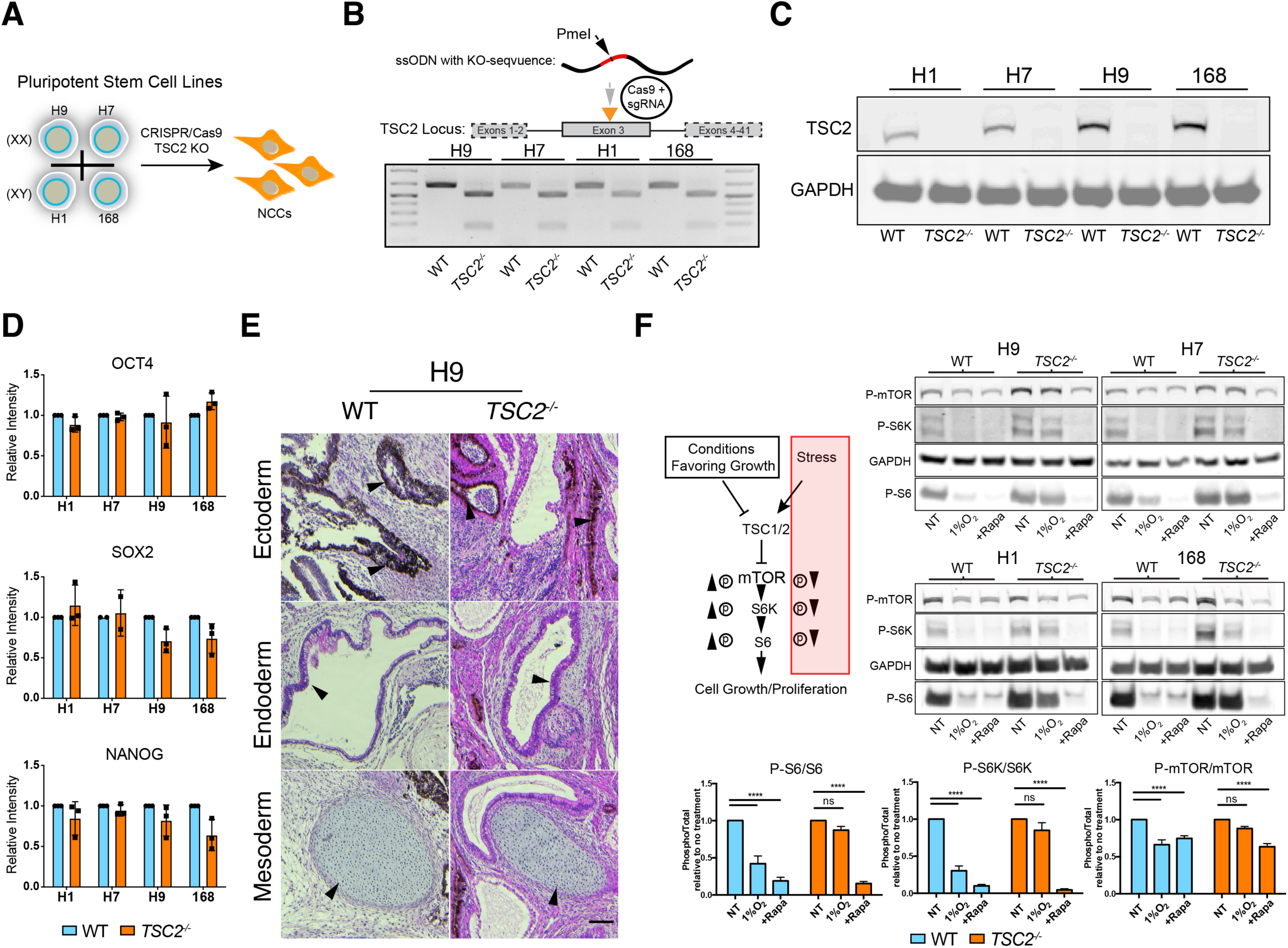
Generation of *TSC2^—/—^* hPSC library. (A) Schematic summary of *TSC2^—/—^* stem cell modeling strategy. (B) Homologous recombination strategy to knockout *TSC2* utilized an ssODN containing a ‘STOP-codon’ donor sequence in all three reading frames containing the unique restriction site, PmeI. PmeI digestion of PCR amplicons containing the target cut site reveals homozygous integration of the donor sequence in *TSC2^—/—^*hPSC lines. (C) Western blot probing for TSC2 in WT and *TSC2^—/—^* hPSC lines demonstrates TSC2 homozygous lines are null. (D) Quantification of fluorescence intensity of pluripotency markers OCT4, SOX2, and NANOG in each hPSC line under maintenance conditions (mean ± SD; n = 12[n = 3 of 168, n = 3 of H9, n = 3 of H7, n = 3 of H1]). No statistical significance between WT and *TSC2^—/—^*samples was observed, confirming normal expression of core pluripotency factors. Values are normalized to respective parental WT hPSC lines. (E) Hematoxylin and eosin (H&E) staining of H9 WT and *TSC2^—/—^* teratoma. Arrows indicate examples of tissues of ectodermal, endodermal, and mesodermal origin, confirming pluripotency. Scale bar, 100 µm. (F) Schematic of expected phosphorylation status of mTORC1 effectors under supportive and stress conditions. Western blots probing phosphorylated mTOR (P-mTOR), phosphorylated S6K (P-S6K), and phosphorylated S6 (P-S6) of samples treated for 6 hours under no treatment, 1% O2, and 100 nM rapamycin conditions demonstrate loss of mTORC1 signaling regulation by *TSC2^—/—^* hPSCs. Densitometry quantification of western blots displaying the ratio of phosphorylated to total protein of mTOR, S6K, and S6 (mean ± SEM; n = 8 [n = 2 of 168, n = 2 of H9, n = 2 of H7, n = 2 of H1]; ∗p < 0.05 by using two-way ANOVA and Tukey’s post hoc analysis).

Gene editing led to the complete ablation of TSC2 (tuberin) protein in the four parental cell lines to generate four isogenic WT and *TSC2^—/—^* pairs of cell lines **(Figure 1C, S1A)**. All hPSC cell lines (WT and *TSC2^—/—^*) can be maintained indefinitely in an undifferentiated state and express hallmark pluripotency markers **(Figure 1D)**. Furthermore, each hPSC line generated ectodermal, mesodermal, and endodermal tissues through *in vivo* teratoma assays **(Figure 1E, S1B)**. The loss of TSC2 function impairs the regulation of mTORC1 signaling, a defining characteristic of TSC lesions. Under normal maintenance conditions, both undifferentiated WT and *TSC2^—/—^* hPSCs exhibited active mTORC1 signaling, as indicated by the phosphorylation of mTOR (Ser2448) and the downstream effectors of the mTORC1 complex, p-S6K (S6K; Thr389) and p-S6 ribosomal protein (S6; Ser235/236). In contrast, upon exposure to stress (6h at 1% O2), WT cells suppressed mTORC1 signaling, while *TSC2^—/—^* hPSCs maintained constitutive phosphorylation of the mTORC1 axis **(Figure 1F, S1C)**. Therefore, all four *TSC2^—/—^* hPSC lines successfully manifest mTORC1 signaling abnormalities observed in TSC-associated lesions and are suitable for studying TSC.

### TSC2^—/—^ NCCs model the PMEL component of LAM

We hypothesized that TSC2 deficient neural crest cells would behave similarly to PMEL^+^ LAM cells. Thus, we employed a twelve-day, attached EB-based dual SMAD inhibition protocol^31^ to differentiate all four sets of hPSCs into NCCs **(Figure 2A)**. During days five through ten of differentiation, cultures exhibited morphological features consistent with NCCs delaminating from the neural tube during embryonic development. The attached EBs formed PAX6^+^ neural rosettes, and the delaminating NCCs expressed key markers for NCC specification (SOX9) and migration (HNK1) (**Figure S2A)**. To enrich for a pure NCC population, selective dissociation was performed on day 12. Regardless of the *TSC2* genotype, this effectively enriched for p75-expressing cells **(Figure S2B-C)**, leading to maintenance cultures that were SOX9^+^/HNK-1^+^ with minimal contamination from PAX6^+^ cells **(Figure 2B).** To further validate neural crest differentiation, we assessed their ability to differentiate into various NCC derivatives. Irrespective of *TSC2* genotype, the stem cell-derived NCCs remain multipotent. They can differentiate into both ectodermal and mesodermal lineages, including SMCs **(Figure S2D-E)**. Finally, differentiation time course RNA-seq analysis of all four isogenic paired lines showed increased expression of neural crest lineage genes such as *SOX9*, *PAX*3, *RXRG, ZEB1, LEF1,* and *NGFR* (encoding p75) compared to the undifferentiated state of NCC **(Figure S2F-H)**. Together, these data demonstrate that TSC2 is not necessary for the differentiation of hPSC into NCCs.

**Figure 2:**
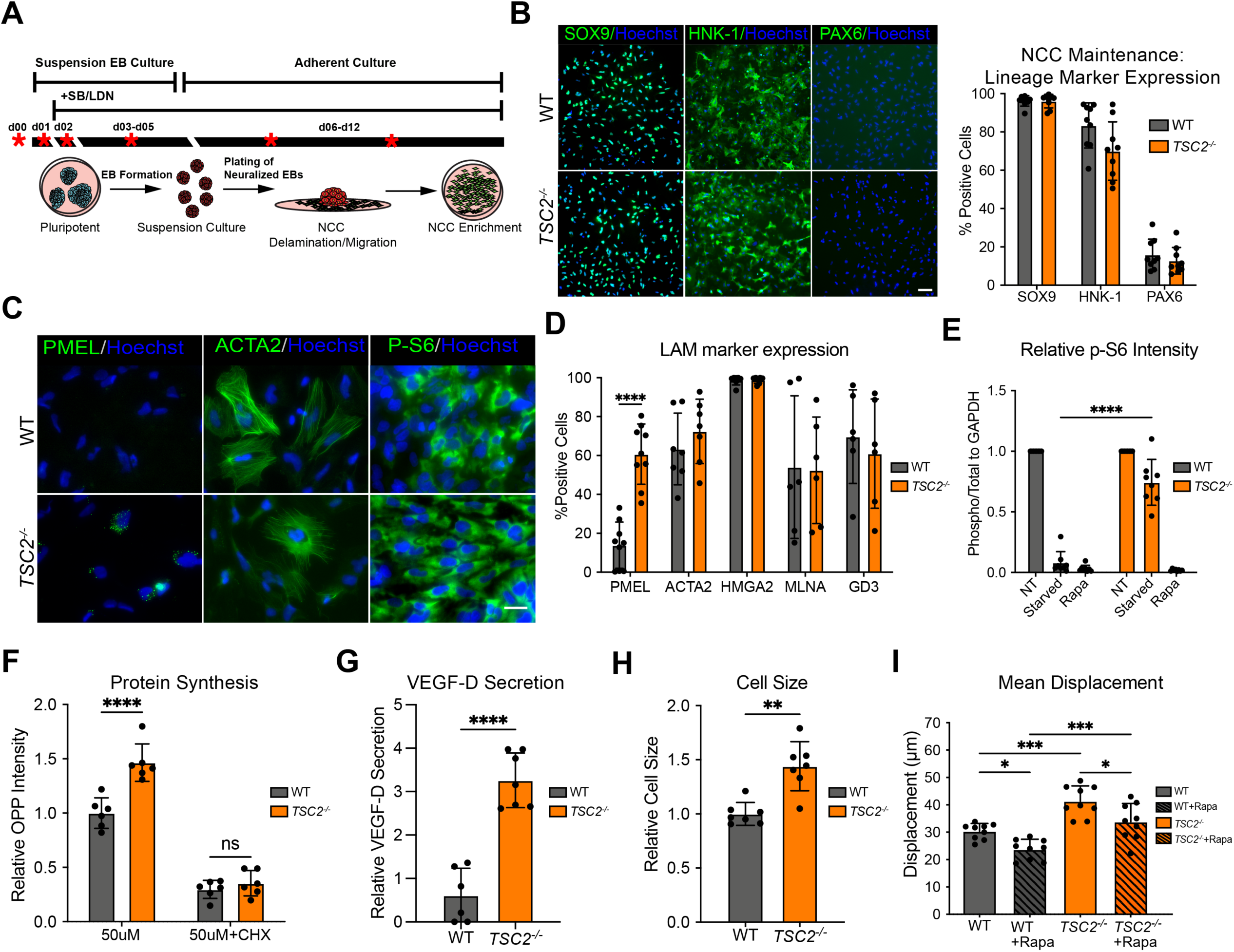
hPSC-derived NCC model key features of pathogenic LAM cells. (A) Schematic representation of EB-NCC differentiation protocols. Red asterisks indicate time points harvested for RNA sequencing. (B) Immunofluorescence staining of passage 1 maintenance NCC cultures, probing for neural crest specific markers, SOX9 and HNK-1, and the neuroepithelial marker, PAX6. Scale bar, 100 µm. Graph shows percentage of NCC populations expressing NCC specific lineage markers SOX9 and HNK-1, and neuroepithelial marker PAX6 (mean ± SD; n = 9 [n = 2 of 168, n = 3 of H9, n = 2 of H7, n = 2 of H1]). No statistical significance was observed between WT and *TSC2^—/—^*NCCs. (C) Immunofluorescence staining of LAM cell marker HMB45, ACTA2 and P-S6. Scale bar, 100 µm. (D) Percentage of WT and *TSC2^—/—^* NCC populations expressing common markers of pathogenic LAM cells (mean ± SD; PMEL [n = 7; 3 of H7, 4 of H9], ACTA2 [n = 7; 3 of H7, 4 of H9], HMGA2 [n=7; 3 of H7, 4 of H9], MLNA [n = 6; 3 of H7, 3 of H9], and GD3 [n = 6; 3 of H7, 3 of H9]; ∗p < 0.05 by using unpaired t-test). (E) Quantification of P-S6 immunofluorescence intensity of NCC maintenance cultures exposed to 24 hours of no treatment, media starvation, or 100 nM rapamycin (mean ± SD; n= 10 [n = 2 of 168, n = 3 of H9, n = 3 of H7, n = 2 of H1] ; ∗p < 0.05 by using two-way ANOVA and Tukey’s post hoc analysis; Values are normalized to no treatment samples within each cell lineage). (F) Protein synthesis levels in WT and *TSC2^—/—^* NCCs with and without cycloheximide (CHX) as determined by O-propargyl-puromycin (OPP) assay, normalized to WT NCCs (mean ± SD; n= 6 [n = 3 of H9, n = 3 of H7]; ∗p < 0.05 by using unpaired t-test; values are normalized to no treatment WT samples). (G) Secretion of VEGF-D in WT and *TSC2^—/—^* NCCs measured via ELISA (mean ± SD; n= 7 [n = 4 of H9, n = 3 of H7]; ∗p < 0.05 by using unpaired t-test; values are normalized to no treatment WT samples). (H) Mean cell size of NCCs under maintenance conditions (mean ± SD; n= 7 [n = 4 of H9, n = 3 of H7]; ∗p < 0.05 by using unpaired t-test; values are normalized to no treatment WT samples). (I) Mean displacement over 6 hours of NCC cultures under maintenance conditions evaluated through time lapse imaging; see also supplemental multimedia (mean ± SD; n= 8 [n = 2 of H9, n = 2 of H7, n = 2 of 168, n = 2 of H1]; ∗p < 0.05 by using one-way ANOVA and Tukey’s post hoc analysis).

We next asked whether the NCC model recapitulates key features of LAM, including the expression of PMEL. Both *TSC2^—/—^* and WT cells expressed common LAM histological markers such as PMEL (HMB45), ACTA2, HMGA2, MLNA (MART-1), and GD3, consistent with the hypothesis that LAM is of neural crest origin^2^. Among these markers, only PMEL was significantly over-expressed in *TSC2^—/—^* NCCs **(Figure 2C-D)**. Additionally, *TSC2^—/—^* NCCs failed to downregulate mTORC1 signaling in response to 24 hours of starvation **(Figure 2E)**, and under normal conditions, *TSC2^—/—^* NCCs exhibited elevated levels of protein synthesis **(Figure 2F)**. We tested whether the ablation of TSC2 in NCCs led to increased secretion of VEGF-D, a critical biomarker used to diagnose LAM. We detected elevated levels of VEGF-D in the supernatant of *TSC2^—/—^*cultures **(Figure 2G)**. Consistent with LAM cell biology, *TSC2^—/—^* NCCs are markedly larger **(Figure 2H)** and show significantly increased migratory potential compared to WT cells, as assessed via live cell tracking over six hours **(Figure 2I, Supplemental multimedia)**. Rapamycin reduced migration in WT and *TSC2^—/—^* NCCs by similar levels, congruent with previous studies demonstrating LAM cell invasion is mTORC1-independent^18,23,30,32^. Together, these data demonstrate that the NCC model exhibits many features reminiscent of LAM lesions.

### TSC2^—/—^ NCCs exhibit transcriptional features of LAM

To gain insight into the initiation of LAM transcriptional landscape, we conducted temporal RNA-seq analysis of *TSC2^—/—^* and WT hPSCs during EB-based NCC differentiation, with time points spanning pluripotency (day 0), EB formation (day 1), neuralization (days 2 and 4), NCC specification (day 7), and NCC enrichment (day 10) **(Figure 2A).** Consistent with prior data **(Figure 1D&E, S1B)**, minimal transcriptional differences were observed at day 0 (19 differentially expressed genes, DEGs) and day 1 (59 DEGs) **(Figure S3A)**. Interestingly, the magnitude of transcriptional dysregulation observed within 24 hours of SMAD inhibition was remarkable. The *TSC2^—/—^* cells exhibited marked dysregulation (189 DEGs on day 2), which intensified by day 4 (2,830 DEGs) and remained elevated at day 10 (609 DEGs) **(Figure S3A)**. To evaluate the biological impact caused by TSC2 deficiency on NCC differentiation, we performed KEGG enrichment analysis on significantly up- and down-regulated DEGs at each time point. From days 0 to 4, *TSC2^—/—^* cells showed consistent upregulation of the “mTOR signaling pathway” and suppression of the “AMPK signaling pathway”, indicating persistent mTORC1 activation. Notably, downregulated genes at day 2 were enriched for the “p53 signalling pathway”, suggesting that TSC2 deficiency disrupts early checkpoints controlling proliferation and apoptosis **(Figure S3B)**.

Given that fully differentiated *TSC2^—/—^*NCCs exhibit multiple LAM-associated features, we next sought to determine whether they also share transcriptional similarities with LAM. Differential expression analysis of *TSC2^—/—^* versus WT NCCs at day 10 identified 609 DEGs at a false discovery rate (FDR) < 0.05, with 207 genes displaying |log_2_FC| > 1 **(Figure S3C).** To gain further biological insight, we performed Gene Ontology (GO) term enrichment on the list of DEGs. Many of the top GO terms identified were also enriched terms in single cell transcriptomic data from primary LAM lesions^24^, such as “Extracellular matrix organization” and “Positive regulation of vascular development” **(Figure S3D)**.

To further evaluate transcriptional similarity to LAM, we compared the LAM cell gene signature derived from scRNA-seq profiling of primary lesions with the protein-coding genes expressed in day 10 *TSC2^—/—^* and WT NCCs. As anticipated, both *TSC2^—/—^* and WT NCCs exhibit substantial overlap with the LAM dataset (111 and 109 genes, respectively), with both gene signatures demonstrating a strong association with LAM as determined by Fisher’s exact test **(Figure S3E).** Notably, the overlapping gene sets from both WT and *TSC2^—/—^*NCCs with LAM include markers ACTA2 (αSMA) and MMP2, as well as matrix-associated proteins implicated in LAM pathology, Lumican (LUM), and Decorin (DCN). Additionally, the LAM gene signature contains many NCC markers, such as *TWIST1*, *SNAI2*, *FRZB*, and *PRRX1*. Importantly, *TSC2*—/— NCCs uniquely expressed critical LAM-associated factors, with VEGFD, PDGFRα, and PDGFRβ specifically enriched at the intersection of the LAM and TSC2—/— gene signatures, reinforcing the relevance of TSC2—/— NCCs as a model of LAM.

### Sexual dimorphic signatures in TSC2^—/—^ NCCs

LAM predominantly affects women, whereas symptomatic LAM is exceedingly rare in men and most often associated with TSC^33,34^. To gain insight into potential sex-related drivers of disease severity, we analyzed our RNA-seq data set to investigate sexual dimorphic transcriptional signatures between WT and *TSC2^—/—^* NCCs. Principal components analysis (PCA) revealed clear separation between male NCC samples (derived from D168 and H1 hPSCs) and female NCC samples (derived from H9 and H7 hPSCs) **(Figure 3A).**

**Figure 3:**
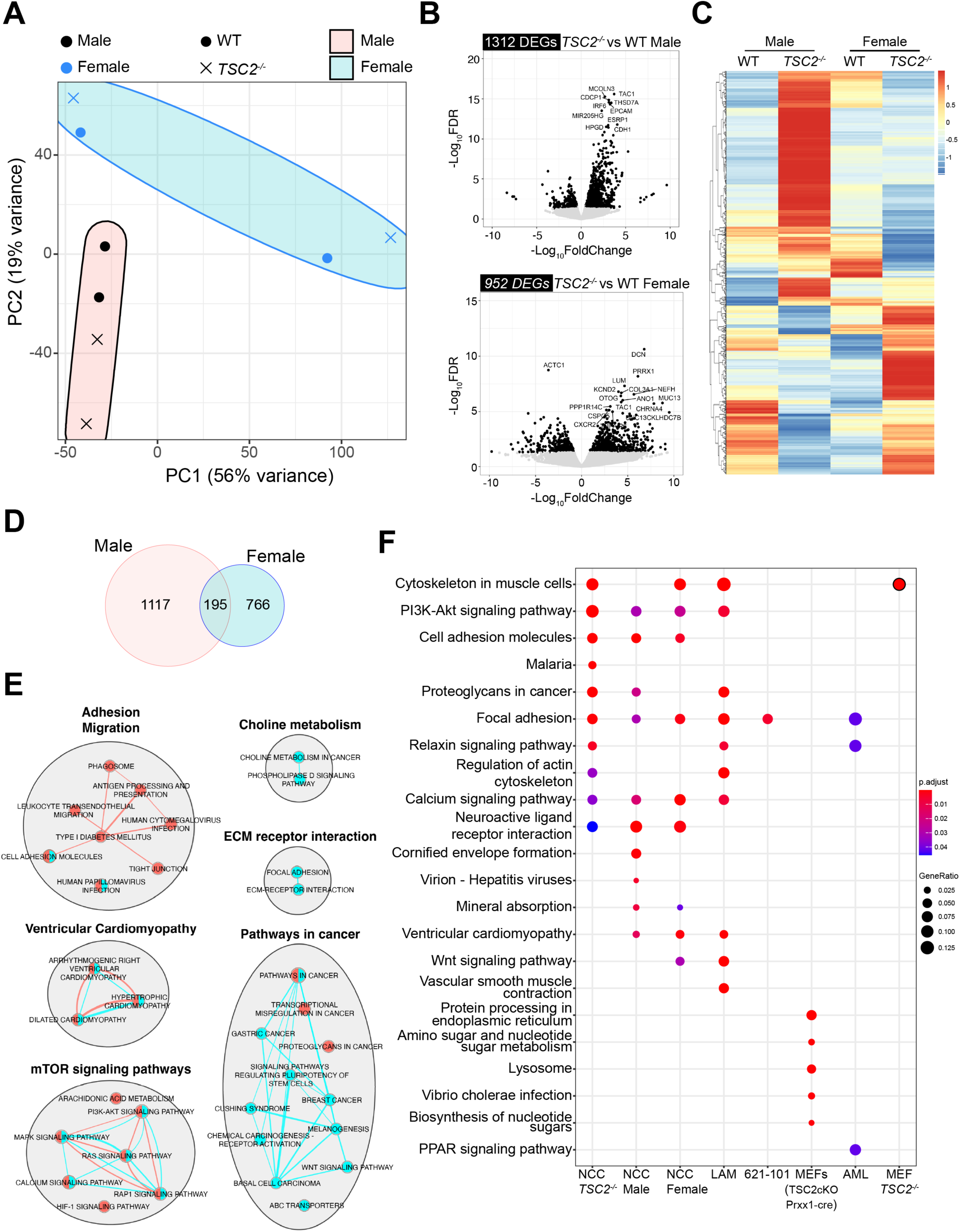
Transcriptomic profiling reveals sex-dependent differences in *TSC2^—/—^*NCCs. (A) Principal component analysis (PCA) of RNA-seq profiles from Day 10 NCCs across all genetic backgrounds (H9, H7, 168, and H1). (B) Volcano plot upon comparing *TSC2^—/—^* vs. WT NCC at day 10 of NCC differentiation. Black points are differentially expressed (FDR < 0.05 and |log2FC| > 1). The 20 most significantly expressed DEGs are noted. Top panel is male bottom is female. (C) Heatmap and hierarchal clustering of differentially expressed genes (DEGs) between male and female NCC lines, and *TSC2^—/—^* and WT samples. (D) Overlap in DEG (FDR < 0.05 and |log2FC| > 1) between male and female gene lists. (E) KEGG pathway network analysis of male and female DEGs (FDR < 0.05 and |log2FC| > 1). Blue nodes correspond to female terms and red to male terms. (F) Comparative KEGG pathway enrichment analysis of *TSC2*^—/—^ NCC, Male *TSC2*^—^/— NCC, Female *TSC2*^—/—^ NCC, LAM cell, 621-101 cell, *Tsc2*^—/—^ NDF, TSC-associated rAML, and *Tsc2*^—/—^ MEF gene signatures. NDF and MEF gene lists were converted to their human orthologs prior to analysis.

We conducted differential expression analysis comparing across genotype (WT vs *TSC2^—/—^*) in male and female lines. At FDR < 0.05, we identified 1328 DEGs (1312 with |log_2_FC|>1) in the male WT versus *TSC2^—/—^* NCCs and 961 DEGs (952 with |log_2_FC|>1) in the female WT versus *TSC2^—/—^* NCCs **(Figure 3B)**. Interestingly, the number of DEGs identified in sex-specific comparisons exceeded those found in the bulk analysis **(Figure S3C),** suggesting that the transcriptional response to loss of TSC2 differs between male and female NCCs. Consistent with this, opposing expression patterns were observed for a subset of genes in male and female *TSC2^—/—^* NCCs **(Figure 3C)**. This canceled out differential signals when sexes were analyzed together. We next examined the overlap between the male and female DEGs. While some overlap was observed (8.6% of the total 2,264 DEGs between male and female), the two data sets were largely distinct **(Figure 3D)**.

To further investigate sexual dimorphic mechanisms, we performed comparative KEGG pathway enrichment analysis. As expected, both male and female DEGs shared enrichment of pathways related to mTORC1 signalling, consistent with the primary role of TSC2 in mTORC1 regulation irrespective of sex **(Figure 3E).** Beyond these shared signatures, male-specific pathways were more broadly distributed across many clusters with notable enrichment for terms linked to “adhesion” and “migration”. In contrast, female-specific pathways were more focused, with a striking enrichment of multiple “Pathways in cancer” terms. This suggests that loss of *TSC2* in female NCCs more strongly promotes oncogenic pathways relevant to LAM.

To benchmark the transcriptional relevance of female *TSC2^—/—^*NCCs as a model for LAM, we compared KEGG pathway enrichment with the LAM gene signature, *TSC2^—/—^* NCCs (bulk, male, and female), along with commonly used LAM cell models, including the immortalized renal angiomyolipoma (rAML) derived 621-101 cell line^35^, *Tsc2*-null mouse embryonic fibroblast (MEFs) lines^36^, and the neonatal dermal fibroblasts (NDFs)^37^ from *Prx1*-Cre-driven conditional *Tsc2*-knockout mice^38^ **(Figure 3F)**. While all *TSC2^—/—^* NCCs shared pathway-level similarity with LAM, female *TSC2^—/—^* NCCs had the greatest number of pathways in common. The unique enrichment of several oncogenic and LAM-relevant pathways in female *TSC2^—/—^* NCCs suggests they more closely recapitulate the transcriptional profile of LAM, establishing them as a relevant human model for LAM.

### A genome-wide CRISPR/Cas9 screen identifies TSC2^−/−^cell vulnerabilities

To identify novel therapeutic targets that could eliminate LAM cells upon inhibition, we performed a genome-wide CRISPR/Cas9-mediated drop-out synthetic lethal screen. To conduct the screen, we first introduced Cas9 into female H9 WT and *TSC2^—/—^* ESCs, which were then differentiated into NCCs **(Figure S4A-B).** To identify synthetic lethal partners for *TSC2,* we employed the genome-wide CRISPR knockout version 2 (GeCKOv.2) lentiviral single guide RNA (sgRNA) library that targets 20,914 genetic elements (19,050 protein coding & 1,864 miRNAs). Following a two-week incubation, changes in sgRNA representation between baseline (day 0) and endpoint (day14) were quantified by deep sequencing, and synthetic lethal genetic elements were ranked using MAGeCK-VISPR and MAGeCKFlute **(Figure S4C).** The screen identified 665 genetic elements that were negatively selected in *TSC2^—/—^* NCCs but not in WT cultures **(Figure 4A)**. The group IV (purple group) data points represent the genetic elements depleted in the *TSC2^—/—^*NCCs and are considered the synthetic lethal gene set.

**Figure 4:**
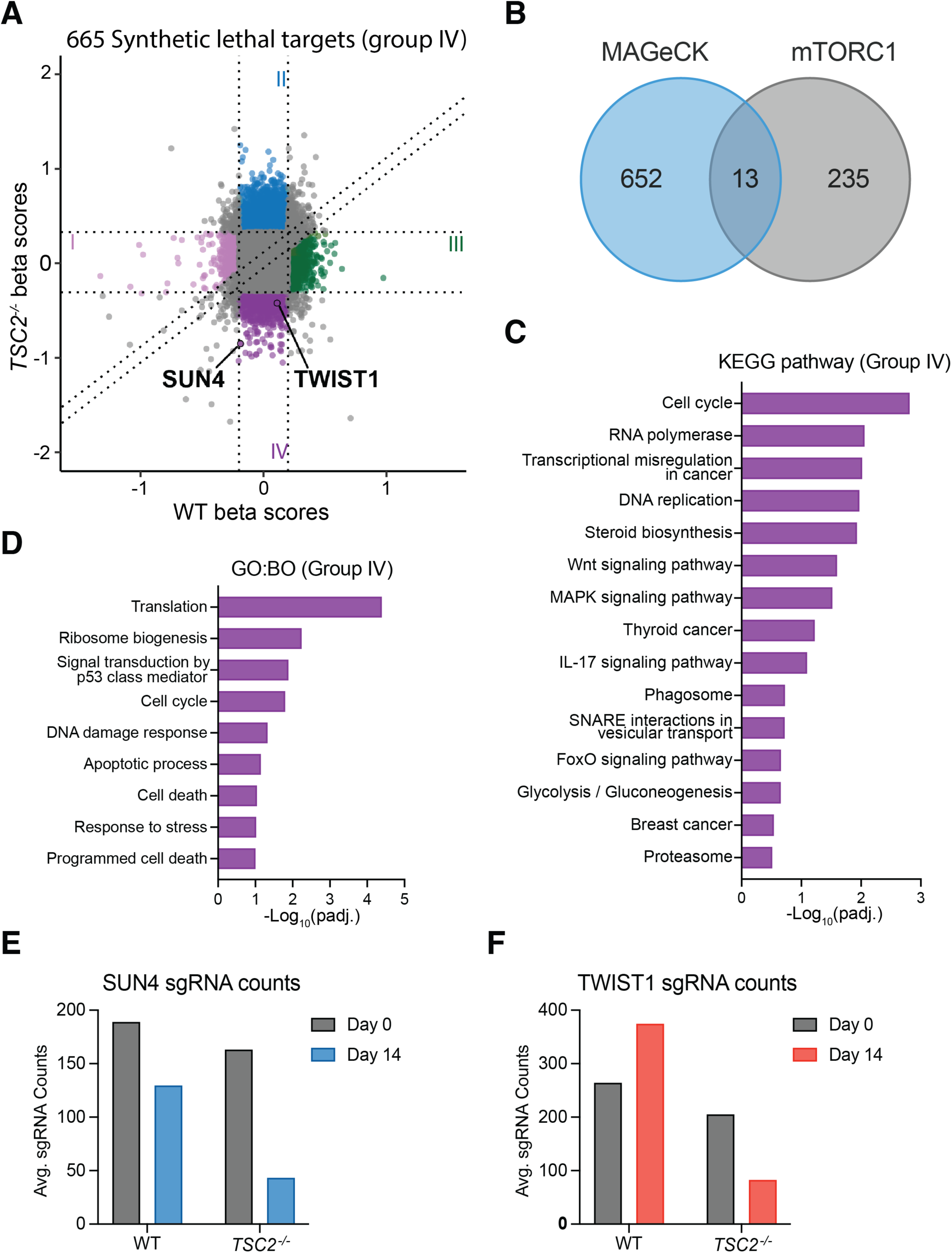
A genome-wide CRISPR/Cas9 screen identifies vulnerabilities in *TSC2^—/—^* NCC. (A) Comparison of gene essentiality scores between WT and *TSC2^—/—^* cells. Positive β-scores indicate genes conferring growth advantage when knocked out, whereas negative β-scores indicate genes required for cell survival or proliferation. Genes were classified into four groups based on their differential essentiality profiles: Group I (pink), genes selectively essential in WT NCCs; Group II (blue), genes whose loss is positively selected in *TSC2^—/—^* NCCs; Group III (green), genes whose loss is positively selected in WT NCCs; and Group IV (purple), genes selectively essential in *TSC2^—/—^* NCCs. (B) Venn diagram of the overlap of group IV synthetic lethal targets and mTORC1 related genes. (C) KEGG pathway enrichment of genes identified as specifically synthetic lethal to *TSC2^—/—^* NCC (group IV). (D) GO Biological processes enrichment of synthetic lethal genes. (E-F) Change in sgRNA counts between day 0 and day 14 for *SUN4* and *TWIST1*.

We next sought to remove gene targets that are associated with mTORC1 from group IV, as it is well recognized that the inhibition of mTORC1 can lead to increased cell survival and would likely be ineffective therapeutic targets in patients undergoing rapamycin treatment. To avoid targets that could be associated with the mTORC1 pathway, we assembled a list of genes associated with the mTORC1 pathway (248 genes) and overlapped it with our group IV *TSC2^—/—^*synthetic lethal targets **(Figure 4B)**. Surprisingly, only 13 genes (1.4%) overlapped between the two gene sets, consistent with the hypothesis that targeting mTORC1-related genes does not cause cell death in *TSC2^—/—^*cells. KEGG pathway gene set enrichment analysis of the remaining group IV genes enriched for terms related to signaling pathways associated with LAM, including the “WNT signaling pathway”, “MAPK signaling pathway”, and “Phagosome” (**Figure 4C).** To gain further biological insight, we conducted GO term enrichment with the remaining group IV gene set (652 genes) **(Figure 4D)**. The most dominant GO biological process terms from this analysis are related to cell death (apoptotic process, signal transduction by p53 mediator, programmed cell death) and the downstream effects of mTORC1 hyperactivation (Translation and ribosome biogenesis). Collectively, these data suggest that the synthetic lethal screen identified non-mTORC1 genes essential to *TSC2*-defficient cells.

Given that removal of mTORC1-associated genes only moderately reduced the size of the group IV gene set, we applied two filtering strategies to further refine the candidate list and reduce the potential of false positives. First, we focused on cross-validating the top group IV genetic elements (protein coding and non-coding RNAs) identified by MAGeCK with those identified by BAGEL, another widely used algorithm for identifying synthetic lethal interactions in CRISPR-based screens **(Figure S4D)**. This comparison revealed a strong overlap of 293 genetic elements with SUN4 emerging as the top target in both data sets. We next compared group IV genetic elements with the RNA-seq DEGs from day 10 WT versus *TSC2^—/—^* NCC differentiation samples, and the LAM-associated genes identified by scRNA-seq of patient lung tissue **(Figure S4E)**. This analysis identified two protein-coding targets, with *TWIST1* being particularly intriguing due to its involvement in various aspects of the metastatic cascade. To validate these findings, we quantified sgRNA read counts at days 0 and 14, which revealed greater depletion of both targets (*SUN4* and *TWIST1*) in *TSC2^—/—^* NCCs than in WT cells **(Figures 4E-F)**. In addition to *TWIST1* and *SUN4* being overexpressed in a range of human cancers, including those not driven by mTORC1 hyperactivation^39–41^, *TWIST1* and *SUN4* are also highly expressed in our *TSC2^—/—^* SMC and renal organoid RNA-seq data sets^23,42^. Together, these observations underscore their potential as broadly relevant therapeutic targets.

### Inhibition of TWIST1 and SUN4 potentiates selective cytotoxicity of TSC2^—/—^ cells independent of rapamycin

The CRISPR-based synthetic lethal screen is based on quantifying sgRNA sequence representation within the population. However, factors beyond cell death, such as inhibition of proliferation, induction of senescence, or loss of adherence could lead to inclusion within the group IV gene set. To validate SUN4 and TWIST1 as synthetic lethal targets for *TSC2^—/—^* cells, we designed two distinct antisense oligonucleotides (ASOs) targeting SUN4 (SUN4_1, SUN4_2) and TWIST1 (TWIST1_1, TWIST1_2). In all *in vitro* experiments, two sets of isogenic female (H9 and H7) WT and *TSC2^—/—^* NCCs were treated with ASOs. To test knockdown efficiency, WT and *TSC2^—/—^* NCCs were treated with SUN4 or TWIST1 ASOs for 48 hours. Following treatment, transcript levels were reduced by >50% **(Figures S5A–B)** and protein abundance decreased by approximately 30—50% relative to control **(Figure 5A-B)**, confirming efficient target suppression. Remarkably, SUN4 and TWIST1 protein levels are approximately two-fold higher in *TSC2^—/—^*NCCs than WT NCCs, suggesting an “oncogene addiction-like” role of these synthetic lethal targets.

**Figure 5:**
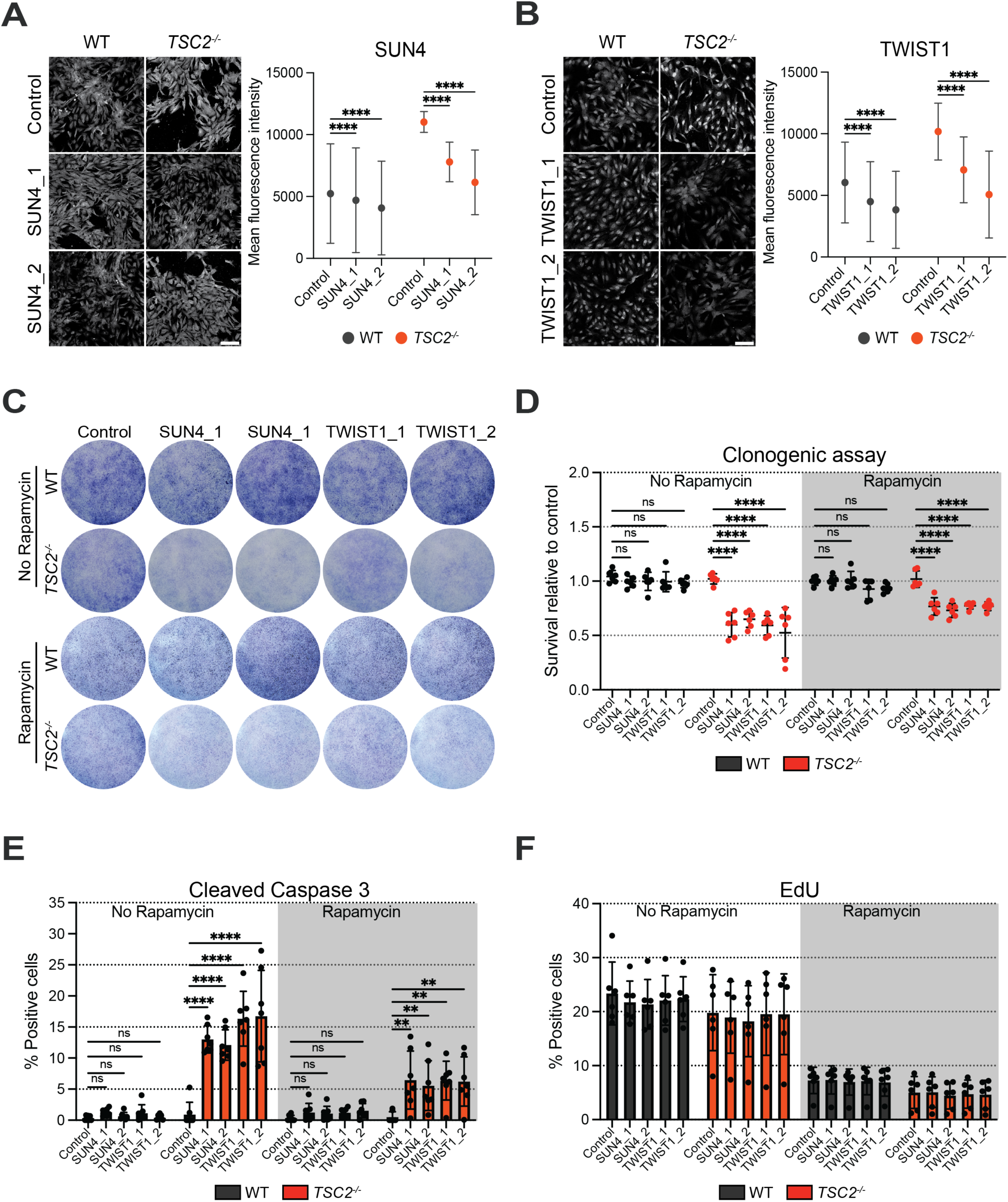
TWIST1 and SUN4 ASOs induce selective cytotoxicity in *TSC2^—/—^*NCCs. (A&B) Representative immunofluorescence images of SUN4 and TWIST1 knockdown after 48 hours of SUN4(SUN4_1, SUN4_2) and TWIST1(TWIST1_1, TWIST1_2) ASO treatment at 500 nM. (A) SUN4-ASO (SUN4_1, SUN4_2). (B)TWIST1-ASO (TWIST1_1, TWIST1_2). Scale bar, 100 µm. Graphs show quantification of fluorescence intensity for knockdown validation of SUN4 (A, right panel) and TWIST1 (B, left panel) in WT and *TSC2^—/—^* NCC (mean ± SD; n= 6 [n = 3 of H9, n = 3 of H7]; ∗p < 0.05 by using two-way ANOVA and Tukey’s post hoc analysis; Values are normalized to Control-ASO within genotype). (C) Clonogenic assay following two ASO treatments and 10 days culture, cells are treated on day 1 and day 6. On day 8 cells are seeded onto 6well plate to assess clonogenicity ±100 nM rapamycin. (D) Clonogenic assay cell counts (mean ± SD; n= 6 [n = 3 of H9, n = 3 of H7]; ∗p < 0.05 by using two-way ANOVA and Tukey’s post hoc analysis; Values are normalized to Control-ASO within genotype). (E) Percentage of cells expressing the apoptosis marker cleaved Caspase 3 in response to 500 nM Control, TWIST1-ASOs or SUN4-ASOs ±100 nM rapamycin after 72 hours. (mean ± SD; n= 6 [n = 3 of H9, n = 3 of H7]; ∗p < 0.05 by using two-way ANOVA and Tukey’s post hoc analysis; Values are normalized to Control-ASO within genotype). (F) Percentage EdU^+^ (proliferating) cells from 3-hour pulse (10μM). NCC were treated with 500 nM of Control-ASO, TWIST1_1-ASO or TWIST1_2-ASO ±100nm rapamycin for 72 hours (mean ± SD; n= 6 [n = 3 of H9, n = 3 of H7]). No statistical significance was observed between Control, SUN4 and TWIST1-ASO treatment groups.

To determine whether SUN4 and TWIST1 knockdown recapitulate the synthetic lethal phenotype identified in the CRISPR screen, clonogenic survival was evaluated following 10 days of ASO treatment in the female *TSC2^—/—^* and WT NCCs. Survival was selectively reduced in *TSC2^—/—^* cells, whereas WT cells exhibited minimal impact on clonogenic growth **(Figure 5C-D)**. Consistent with impaired survival, approximately 15% of *TSC2^—/—^* NCCs were positive for the apoptosis marker cleaved caspase-3 (CASP3) after 72 hours of TWIST1 or SUN4 ASO treatment compared to minimal apoptosis in WT cells **(Figure 5E)**. Sytox incorporation further supported selective cytotoxicity in *TSC2^—/—^*cells **(Figure S5C)**. Consistent with increased cell death, micronuclei frequency was elevated in *TSC2^—/—^* cells and metabolic activity was decreased following ASO treatment **(Figure S5D-E)**. Notably, rapamycin partially reduced the cytotoxic response **(Figure 5E)**. However, ASO treatment alone did not impact proliferation, whereas rapamycin markedly reduced EdU incorporation in both WT and *TSC2^—/—^* NCCs **(Figure 5F)**, suggesting that slowed proliferation likely accounts for the rapamycin-dependent decrease of cytotoxicity.

### RNA-seq analysis reveals anti-metastatic effects upon knockdown of TWIST1 and SUN4 in TSC2^—/—^ cells

We next aimed to elucidate the mechanisms of action using bulk RNA-seq on WT and *TSC2^—/—^* NCCs treated with control, SUN4, or TWIST1 ASOs **(Figure S6A).** Principle component analysis (PCA) revealed genotype as the primary driver of transcriptional variation, with treatment effects contributing to the secondary axis, driven largely by SUN4 and TWIST1 ASO treated *TSC2^—/—^* samples **(Figure 6A)**. To explore genotype selective changes induced by SUN4-ASO and TWIST1-ASOs, we first utilized DESeq2 for differential expression analysis. We compared across genotypes (*TSC2^—/—^* vs WT) and treatments (Control-ASO, SUN4-ASO, TWIST1-ASO), applying an FDR < 0.05. This analysis identified 1705 DEGs (787 with |Log_2_FC| > 1) in *TSC2^—/—^*compared to WT NCCs treated with Control-ASO. SUN4-ASO treatment resulted in 2902 DEGs (940 with |Log_2_FC| > 1), while TWIST1 knockdown with TWIST1-ASO revealed 2206 DEGs (787 with |Log_2_FC| > 1) **(Figure S6B-D).**

**Figure 6.**
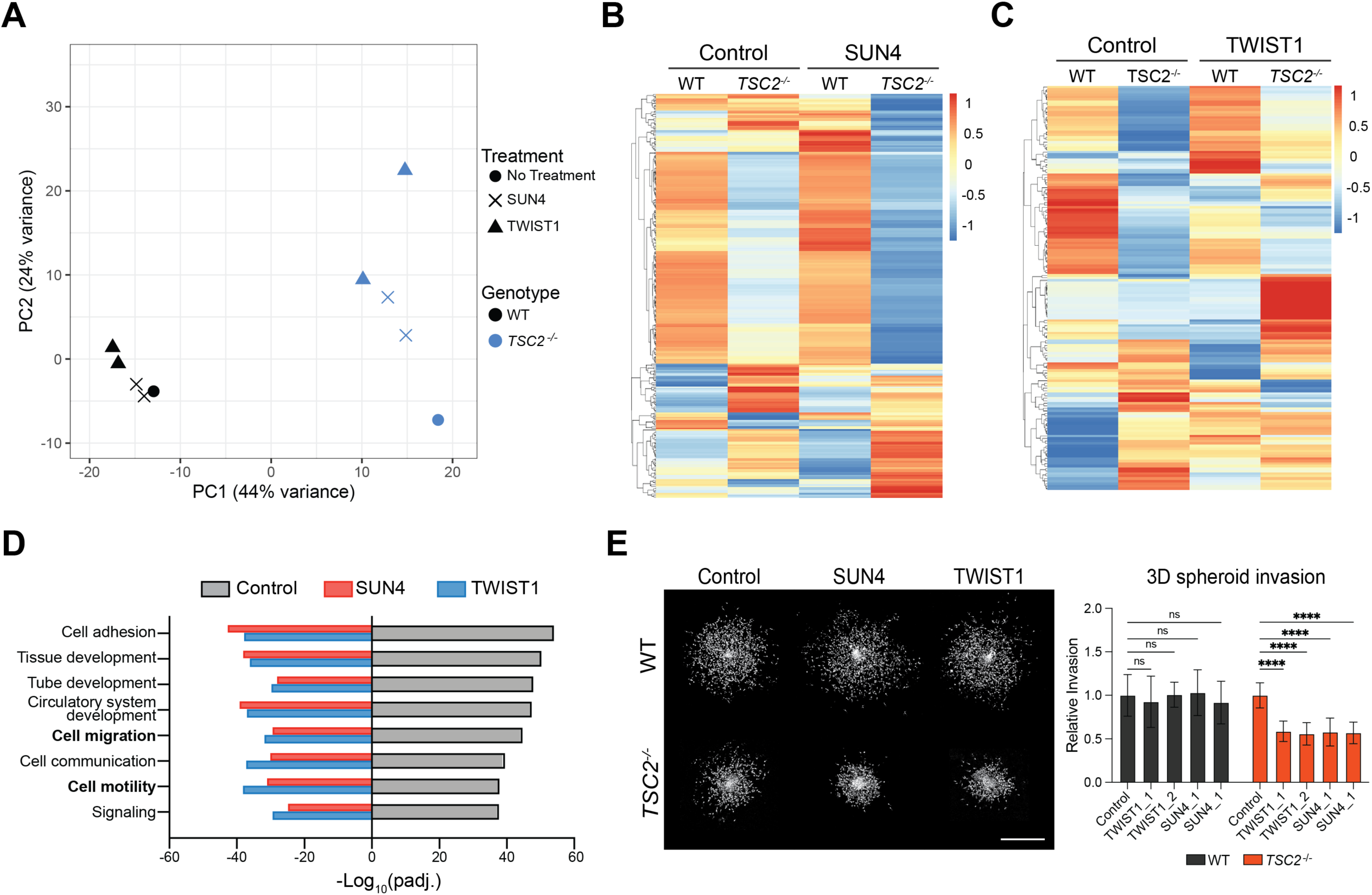
Inhibition of TWIST1 and SUN4 with respective ASOs reverse metastatic effects of *TSC2^—/—^* cells. (A) Principal components analysis (PCA) of bulk RNA-seq samples. (B&C) Heat map and hierarchal clustering of DEGs found significant (FDR < 0.05) in the interaction between genotype and treatment (B, SUN4 261 genes, C, TWIST1 161 genes). (D) Bar plots of GO Biological process enrichment analysis comparing DEG lists (FDR < 0.05 and log2FC > 1) for Control-ASO treated *TSC2^—/—^* vs WT cells and downregulated interaction-term DEGs (FDR < 0.05 and log2FC < -1) specifically affecting *TSC2^—/—^* cells. The top eight most significant terms that are common among all three groups are shown. (E) Fluorescence visualization of WT and *TSC2^—/—^* NCC cell invasion into Matrigel after three days of treatment with 250 nM Control-ASO, TWIST1-ASO or SUN4-ASO. Image of single Z plane. Scale bar, 500 μm (Left panel). Right, relative invasion of WT and *TSC2^—/—^* NCC into the surrounding matrix quantified as area covered by invading cells relative to the core spheroid (mean ± SD; n= 6 [n = 3 of H9, n = 3 of H7]; ∗p < 0.05 by using two-way ANOVA and Tukey’s post hoc analysis; Values are normalized to Control-ASO within genotype).

To identify genes whose transcriptional response to SUN4-ASO or TWIST1-ASO treatment differed between WT and *TSC2^—/—^* cells, rather than those reflecting baseline genotype differences alone, we examined the genotype:treatment interaction term in the DESeq2 model. We identified 261 and 161 DEGs (FDR < 0.05) for SUN4-ASO and TWIST1-ASO, respectively. Hierarchical clustering of these genes across all conditions (WT and *TSC2^—/—^*± ASO-treatment) highlighted transcriptional responses specific to the treated *TSC2^—/—^* samples **(Figure 6B-C).** GO term enrichment analysis of genes showing treatment-dependent expression changes restricted to the *TSC2^—/—^* condition demonstrated significant upregulation of terms clustering into categories related to “Apoptotic processes” and “Programmed cell death”, alongside downregulation of term clusters linked to metastatic processes, including “Cell adhesion” and “Motility, locomotion, migration.” **(Figure S6E-F)**. Strikingly, many of the pathways suppressed by SUN4-ASO and TWIST1-ASO treatment were conversely upregulated in Control-ASO treated *TSC2^—/—^*versus WT **(Figure 6D),** indicating that SUN4-ASO and TWIST1-ASO treatments disrupt transcriptional programs associated with LAM. While SUN4-ASO and TWIST1-ASO treatment converge on shared cell death and anti-metastatic signatures, their regulatory profiles diverged. SUN4, a component of the linker of nucleoskeleton and cytoskeleton (LINC) complex^43,44^, preferentially enriched for processes related to “Plasma membrane organization” **(Figure S6E)**, whereas TWIST1, a pioneer transcription factor^45–47^, enriched for terms associated with “Epigenetic regulation” **(Figure S6F)**. Together, these data indicate that although both ASOs induce overlapping cytotoxic and anti-metastatic transcriptional programs in *TSC2^—/—^*cells, they do so through distinct mechanistic processes.

To validate the RNA-seq findings and assess the efficacy of TWIST1-ASO and SUN4-ASO in reducing invasion through the extracellular matrix, we employed a 3D spheroid system in which cell spheroids were embedded in Matrigel. Briefly, *TSC2^—/—^* and WT NCC spheroids were pre-formed by incubating 1,000 cells per spheroid in ultra-low attachment plates overnight, followed by embedding in Matrigel and monitoring their invasion into the surrounding matrix **(Figure 6E)**. Notably, TWIST1 and SUN4 ASO treatment resulted in a pronounced anti-invasive effect in *TSC2^—/—^*NCCs, with minimal impact on WT NCCs **(Figure 6E)**. Collectively, these data indicate that TWIST1 and SUN4 are critical for the invasive capacity of *TSC2^—/—^* NCCs, and ASO-mediated knockdown of these genes serves as an effective strategy to inhibit invasion.

### TWIST1-ASO and SUN4-ASO treatments drive sustained tumor regression beyond rapamycin in vivo

We next sought to evaluate the *in vivo* efficacy of TWIST1 and SUN4 ASOs. Previous reports demonstrated that the loss of TSC2 in our cell model alone was insufficient to drive tumorigenicity in subcutaneous xenotransplantation assays^23^. To address this limitation, we turned to the gold standard TSC animal tumor model, an allograft model that utilizes *Tsc2^—/—^* cells (known as 105k) derived from renal cell cystadenoma that spontaneously developed in a *Tsc2^+/—^C57Bl/6* mouse^48^. 105k cells were injected subcutaneously into immune competent (*C57Bl/6*) mice and allowed to grow to 100mm^3^ before administering treatment. *In vivo* ASO dosing strategies vary substantially depending on route of administration and disease context^49,50^. Thus, we first sought to evaluate the antitumor activity of TWIST1-ASOs and SUN4-ASOs *in vivo* using a short-term low-dose efficacy study comparing to standard of care rapamycin in the allograft model. Tumor bearing mice were treated intraperitoneally with vehicle, rapamycin (1.14 mg/kg), SUN4-ASOs (1.12 mg/kg), or TWIST1-ASOs (1.12 mg/kg) for ten days **(Figure S7A)**. Under these short-term, low-dose conditions, SUN4-ASO and TWIST1-ASO treatment induced tumor regression to a comparable degree to rapamycin, with no clear separation between ASO groups and rapamycin **(Figure S7B-D).** These results establish that ASO based treatments exhibit measurable *in vivo* activity comparable to rapamycin even under conservative dosing conditions.

To elicit maximal antitumor responses with ASOs, we surmised that prolonged exposure and higher cumulative dosing may be required. Given that short-term treatment at reduced ASO doses produced antitumor effects comparable to rapamycin, we next asked whether differences between ASO and rapamycin treated tumors would emerge under extended treatment conditions. We further sought to determine whether ASO treatments could induce sustained tumor regression following treatment withdrawal, as rapamycin has been shown to permit rapid tumor regrowth after cessation of therapy^51^. We therefore implemented a “test–withdrawal–monitor” study design similar to that used in LAM and TSC clinical trials^17,18^ consisting of an extended treatment regimen of 30 days, followed by a 30-day treatment withdrawal period, to assess both maximal tumor response and durability after cessation of therapy **(Figure 7A)**.

**Figure 7.**
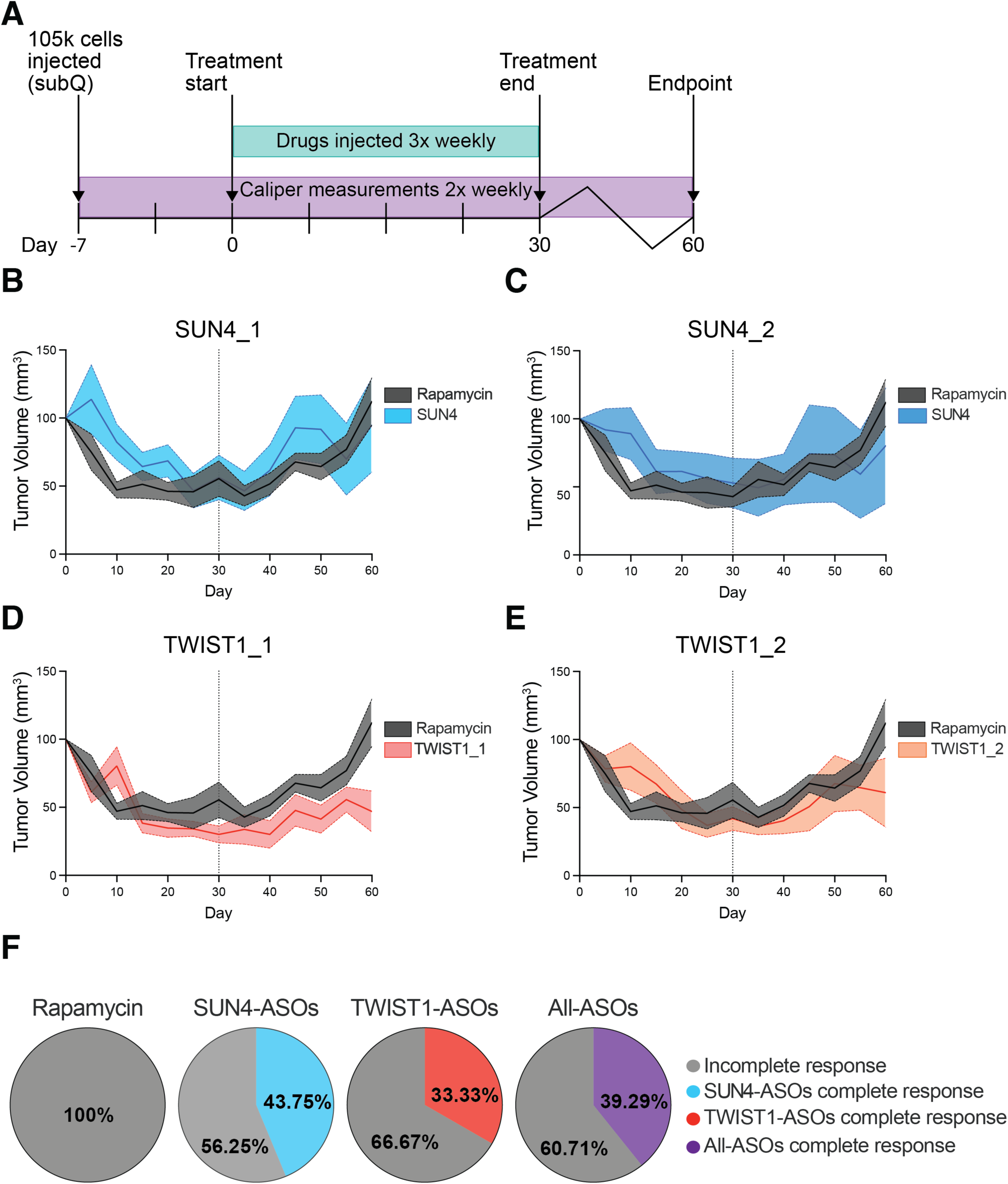
SUN4-ASO and TWIST1-ASO treatment induces complete regression in a subset of tumors *in vivo*. (A) Schematic diagram of treatment strategy. 105k cells were injected subcutaneously, and after the tumor volumes reached 100 mm^3^ (2 tumors per mouse), mice were divided into the following treatment groups: Rapamycin (1.14 mg/kg, n = 4), SUN4_1 (10 mg/kg, n = 4), SUN4_2 (10 mg/kg, n = 4), TWIST1_1 (2.25 mg/kg, n = 3), TWIST1_2 (2.25 mg/kg, n = 3). Animals were treated 3 times a week via i.p. injections. (B-E) Plots of tumor volumes during 4 weeks of treatment followed by 30-day treatment withdrawal. Solid lines represent mean tumor volume for each group, and shaded regions denote SEM. The vertical dash line indicates the time point of treatment cessation (B: SUN4_1, C: SUN4_2, D: TWIST1_1, E: TWIST1_2). (F) Pie charts illustrating the proportions of tumors exhibiting complete regression versus residual tumors across treatment groups following withdrawal.

Under the “test–withdrawal–monitor” study design, SUN4-ASOs (10 mg/kg) and TWIST1-ASOs (2.25 mg/kg) again elicited marked tumor response by the end of the 30-day dosing period, comparable to the response observed with rapamycin (1.14 mg/kg) **(Figure 7B-E)**. Notably, complete tumor regression was observed in a subset of ASO-treated tumors **(Figure S7E**). Following treatment withdrawal, ASO and rapamycin-treated cohorts exhibited regrowth. However, a subset of ASO-treated tumors displayed complete remission, defined by the absence of detectable tumors at necropsy. In contrast, all rapamycin-treated tumors uniformly retained residual disease and demonstrated tumor regrowth following treatment cessation **(Figure 7F, S7E-F)**. Together, these data demonstrate that while TWIST1-ASOs, SUN4-ASOs and rapamycin treatments show comparable short-term antitumor activity, extended ASO treatment uniquely eradicated a subset of tumors that persists following treatment withdrawal.

## Discussion

Experimental models that faithfully recapitulate the molecular and phenotypic features of LAM remain limited, hindering efforts to identify new therapeutics. Here, we establish a human stem cell-based platform designed to model the PMEL-expressing cellular component of LAM. Using this system, we demonstrate that *TSC2^—/—^* NCCs recapitulated numerous pathognomonic features of LAM cells, including PMEL and ACTA2 expression, VEGF-D secretion, and dysregulated mTORC1 signaling. A genome-wide CRISPR dropout screen identified *SUN4* and *TWIST1* as synthetic lethal targets of *TSC2^—/—^* cells. Target suppression using ASOs induced selective cytotoxicity *in vitro* and eradicated tumors in a subset of mice in a test-withdrawal-monitor study design, revealing therapeutic vulnerabilities in *TSC2^—/—^* cells beyond mTORC1 inhibition.

The cell of origin of LAM has been in question for decades, with numerous candidates proposed^16,24,52–54^, including NCCs^2,55^, which may provide a useful context in modeling LAM-like phenotypes. LAM lesions exhibit a distinctive combination of melanocytic and smooth muscle features^2,16^, both of which are derivatives of NCCs^2,56^. As expected, both *WT* and *TSC2^—/—^* NCCs express common markers of LAM such as ACTA2, MELNA, GD3, and HMGA2. This supports that the neural crest lineage itself is a representative cell type for LAM. However, *TSC2^—/—^* NCCs specifically display increased abundance of pathognomonic LAM markers, PMEL and VEGF-D, while exhibiting classic oncogenic phenotypes of mTORC1 hyperactivation. Although our data do not resolve the cellular origin of LAM, they demonstrate that NCCs possess broad modeling potential since it can generate multiple cell types present in LAM nodules and capture key molecular and transcriptional features of LAM cells.

LAM occurs almost exclusively in women; a substantial body of clinical and experimental evidence implicates estrogen signaling as an important contributor to disease progression^57–68^. However, limited efficacy of hormone-targeting therapies suggests that biological sex may influence disease pathogenesis through mechanism that extend beyond estrogen signaling alone^69–71^. Despite this possibility, relatively little work has explored mechanisms by which biological sex may influence disease progression. This gap is partly due to the rarity of symptomatic LAM in males and the limited availability and expansion capacity of primary LAM and angiomyolipoma samples^28,33,34^. In this context, the isogenic stem cell platform described here provides a system for which to investigate sex-dependent responses to loss of TSC2. While male and female *TSC2^—/—^* NCCs exhibit strong enrichment for mTORC1 pathway signatures and share transcriptional similarities with patient-derived LAM cells, female *TSC2^—/—^* NCCs enriched for melanogenesis and cancer-associated transcriptional programs. These findings indicate that while mTORC1 dysregulation represents a central molecular feature of LAM, cell-intrinsic sex-dependent biological factors may influence how cells respond to loss of TSC2 and shape lineage-associated transcriptional programs, potentially contributing to the strong female predominance observed in LAM.

LAM is a rare disease, and like many other rare disorders, therapeutic discovery is challenging. The only disease-modifying therapy currently available is rapamycin, which slows lung function decline and reduces disease-associated adverse events^72–75^. However, its effects are largely cytostatic, and lesions rebound if treatment is withdrawn^17,75,76^. These observations suggest that additional pathways sustain the survival of LAM cells and may represent opportunities for therapeutic intervention. Consistent with this supposition, our genome-wide CRISPR dropout screen revealed that known canonical components of the mTORC1 signaling pathway were underrepresented among the identified *TSC2^—/—^*synthetic lethal targets. Notably, the two targets selected, *TWIST1* and *SUN4* are not established members of the mTORC1 pathway. TWIST1 is best known for its role in regulating EMT^47,77,78^. Previous studies have demonstrated cross-talk between TWIST1-driven EMT programs and mTORC1 signaling pathways but no studies have definitively shown TWIST1 to be a canonical mTORC1 marker^79–81^. Proteins of the LINC complex, including SUN4, physically couple the cytoskeleton to the nuclear envelope and transmit mechanical forces to the nucleus^82^. Through these interactions, nuclear-cytoskeletal connections regulate pathways that influence cellular growth signaling networks that may converge on mTORC1 signaling^83–85^. Together, these observations suggest that TSC2-deficient cells rely on previously unrecognized dependencies that extend beyond canonical mTORC1 signaling for survival.

ASOs have emerged as a powerful therapeutic platform with demonstrated clinical utility across a range of diseases, including cancer^86^. Notably, ASO-based therapies offer significant potential for the treatment of rare diseases, where limited patient populations can make conventional drug development challenging^87^. These therapies provide a flexible and scalable strategy for precisely targeting genes of interest, with approaches that can be adapted to small patient cohorts or even individual patients^87–91^. The growing number of ASO-based therapeutics that have received regulatory approval or are currently under clinical investigation further underscores the translational feasibility of this approach and its potential relevance for LAM^92^. Within this context, TWIST1 and SUN4 represent promising therapeutic targets. Inhibition of these targets disrupt pathways required for the survival of *TSC2^—/—^*cells, indicating that targeting these genes enables cytoablative therapeutic strategies capable of eliminating TSC2-deficient cells rather than simply suppressing disease progression.

Targeting of TWIST1 and SUN4 with ASOs may have implications beyond LAM. TWIST1 has long been recognized as a driver of tumor progression in multiple cancer types^39,40^, yet transcription factors have historically been difficult to target therapeutically, in part because they function through protein–protein and protein–DNA interactions rather than defined ligand-binding sites that are amenable to small-molecule inhibition^93,94^. SUN4 is less well characterized and currently lacks targeted therapeutic approaches, but has been linked to cancer progression^41,43^ and, as a component of the LINC complex, may contribute to pathways that support tumor growth and invasion^95–97^. Our findings suggest that ASO-based approaches provide a strategy to directly suppress these factors, and advances in delivery platforms, including lipid nanoparticles^98,99^ ,offer opportunities for tissue-specific targeting in cancers. Together, these observations suggest that ASO-mediated targeting of TWIST1 and SUN4 may have broader therapeutic relevance across multiple disease contexts.

Treatment with TWIST1 or SUN4-targeting ASOs produced durable anti-tumor responses *in vivo* and, notably, resulted in complete tumor eradication in a subset of animals in a test-withdrawal-monitor study. The persistence of the cytoablative responses after treatment withdrawal suggests that inhibition of these targets can eradicate LAM/TSC cells rather than simply suppressing lesion growth. This contrasts with the effects typically observed with mTOR-directed therapies for LAM, which largely control disease during treatment but rarely produce durable regression following therapy cessation^17,75,76^. The variability in tumor responses suggests variable delivery or uptake of ASOs within tumors may limit the extent of target inhibition *in vivo*, and some tumors may require longer treatment durations, higher dosing, or delivery via lipid nanoparticles to achieve sufficient target suppression. Together, it highlights the need to better understand the factors that determine sensitivity to TWIST1 or SUN4 inhibition *in vivo*.

In conclusion, this study establishes that human stem cell derived *TSC2^—/—^* NCCs serve as an accurate cell model for LAM cells that demonstrates key molecular and phenotypic features of the PMEL-expressing LAM cell population. Through transcriptional profiling of NCCs, we uncovered previously unappreciated sex specific differences that could drive disease incidence in women and should be further explored. Furthermore, the combination of the CRISPR screen and the NCC model provides a platform to identify therapeutic vulnerabilities associated with loss of TSC2. Our screen serves as a valuable resource for the LAM community because it nominates genes that, upon loss of function, may cause selective cytotoxicity in LAM cells. We show here, using two examples (TWIST1 and SUN4), that our screen identified targets whose inhibition can selectively impair the survival of *TSC2^—/—^* cells. This validation supports the broader utility of the screen and suggests that additional candidates may represent actionable vulnerabilities in LAM.

## Limitations

There are limitations of this study that should be considered. Although the human stem cell–derived NCC model recapitulates key molecular and phenotypic features of LAM, it does not fully capture the complexity of the cellular and microenvironmental context present in patient lesions. In addition, the *in vivo* preclinical experiments were performed using the 105K allograft model, which is considered the gold standard TSC tumor model. While this model enabled evaluation of anti-tumor activity *in vivo*, LAM cells are non-transformed and demonstrate a slower proliferation rate than *Tsc2^—/—^*cells derived from renal cell cystadenoma that spontaneously developed in a *Tsc2^+/—^ C57Bl/6* mouse ^48^ . Furthermore, although TWIST1 and SUN4 inhibition produced selective cytotoxic effects in *TSC2^—/—^* cells, the precise molecular mechanisms underlying these responses remain to be fully defined. Future studies addressing these questions and evaluating optimized ASO delivery strategies will be important for advancing TWIST1 and SUN4-directed therapies toward clinical application.

## Acknowledgements

The authors would like to acknowledge the assistance of the Ottawa Hospital Research Institute (OHRI) Flow Cytometry and Cell Sorting Core Facility (RRID:SCR_023349), the Ottawa Human Pluripotent Stem Cell Facility (RRID:SCR_027437), the OHRI Bioinformatics Core Facility (RRID:SCR_022466), OHRI Stem Core (RRID:SCR_012601), and the OHRI High Content Imaging Core. We acknowledge the generous funding was provided to WLS via grants from the Canadian Institutes of Health Research (CIHR) [FRN-153188, PX2-204176], Natural Sciences and Engineering Research Council of Canada (NSERC) [RGPIN-2022-05138], and The LAM Foundation [LAM0136P01-19] to WLS. We also recognize Catherine Lawrence and Laugh Out LAM for their inspiration and financial support related to LAM and TSC research. WLS has been supported by the University of Ottawa Distinguished Research Chair in Disease Modeling and Therapeutic Discovery and a Tier 1 Canada Research Chair in Integrative Stem Cell Biology.

## Declarations of interest

WLS is a member of the Scientific Advisory Board of The LAM Foundation and Scientific Director of the Ottawa Human Pluripotent Stem Cell Facility and OHRI High Content Imaging Core.

## Methods and Materials

### RESOURCE AVAILABILITY

#### Lead Contact

Further information and requests for resources and reagents should be directed to and will be fulfilled by the lead contact, William L. Stanford

#### Materials Availability

The cell lines (*TSC2^—/—^* hPSCs) and plasmids generated in this study are available upon request to the lead contact.

#### Data Availability

RNA-seq data generated in this paper has been deposited at GEO (accession number GSE137614, GSE338669) and are publicly available as of the publication date. This paper analyzes existing, publicly available data. Sources for these datasets are listed in the key resources table. All other data are available upon request to the lead contact.

### EXPERIMENTAL MODEL DETAILS

#### Human Pluripotent Stem Cells

hESC lines H9 (WiCell WA-09, female), H7 (WiCell WA-07, female), H1 (WiCell WA-01, male) and the male induced pluripotent stem cell line 168 (Chen et al., 2017), as well as their respective *TSC2^—/—^* derivatives, were maintained on diluted Matrigel at 0.16 mg/mL (BD Biosciences #354230). Cells maintained in either Essential 8 (E8) or mTESR Plus media (Stem Cell Tech #100-0276) at 37 °C, 5 % O_2_, 10 % CO_2_. Media was replaced daily, and hPSCs were passaged in aggregates every 4-5 days, before cells reached 80% confluency, using 0.5 mM EDTA. Spontaneously differentiated cells are manually removed prior to passing.

E8 media recipe: DMEM/F-12 (Thermo Fisher #11330), 64 µg/mL Ascorbic Acid 2 Phosphate Mg (Sigma #A8960), 14 ng/mL sodium selenium (Sigma #S526), 0.1 µg/mL FGF2 (Thermo Fisher #PHG0263), 19.4 µg/mL human Insulin (Wisent #511-016-CM), 10.7 µg/mL transferrin (Sigma #T0665), 2 ng/mL TGF-β1 (Thermo Fisher #PHG9202), 543 µg/mL NaHCO3 (Sigma #5761) (Chen et al., 2011).

#### 105K cells

105K cells gift from the lab of Dr. Elizabeth Henske. The cells were derived from a *Tsc2^+/—^ C57Bl/6* mouse renal tumor and isolated in the laboratory of Dr. Henske. These cells were confirmed to have loss of the second allele of *Tsc2* by PCR, loss of tuberin expression, and increased phospho-S6 levels by immunoblotting ^43^. Cells were cultured in Dulbecco’s Modified Eagle Medium (DMEM) supplemented with 10 % FBS, 100 μg/mL of penicillin, and 100 μg/mL of streptomycin.

#### Mouse teratoma formation

Teratoma assays (see Teratoma Assays in method details section below) in 6–8-week-old female *NOD.Cg-Prkdc^scid^ Il2rg^tm1Wjl^/SzJ* (*NSG*; Jackson Labs, #005557) mice. Experiments were conducted at the University of Ottawa Animal Care Vivarium in cooperation with University of Ottawa Animal Care and Veterinary Services (ACVS) in accordance with the Canadian Council on Animal Care Standards and the Province of Ontario’s Animals for Research Act. *NSG* mice used in this study were healthy, immunocompromised and maintained in sterile housing conditions. *NSG* mice were not involved in any previous procedures or tests prior to initiation of teratoma assays. *NSG* mice were monitored daily by the ACVS; mice were euthanized during an experiment if indications of ill-health or discomfort were detected (e.g. weight loss, hunched posture, lethargy, rough coat).

#### 105K allograft

Allograft assay (see allograft assay in method details section below) in 7–20-week-old male and female *C57BL/6* (Jackson Labs, #000664) mice. Experiments were conducted at the University of Ottawa Animal Care Vivarium in cooperation with University of Ottawa Animal Care and Veterinary Services (ACVS) in accordance with the Canadian Council on Animal Care Standards and the Province of Ontario’s Animals for Research Act. *C57BL/6* mice used in this study were healthy and immunocompetent were maintained in sterile housing conditions. *C57BL/6* mice were not involved in any previous procedures or tests prior to initiation of allograft assays. *C57BL/6* mice were monitored daily by the ACVS; mice were euthanized during an experiment if indications of ill-health or discomfort were detected (e.g. weight loss, hunched posture, lethargy, rough coat).

### METHOD DETAILS

#### NCC Cell Culture

NCCs were maintained on a thin layer (0.16 mg/mL) Matrigel coated tissue culture treated plates (TC plates). NCCs were maintained in neural induction media (NIM): 1:1 DMEM/F-12: Neural Basal Media (Thermo Fisher #21103-049), 0.5 x N2 Supplement (Thermo Fisher #17502-048), 0.5 x B27 Supplement (Thermo Fisher #17504-044), 5 µg/mL human Insulin, 0.02 µg/mL FGF2, 0.02 µg/ml hEGF), 0.5 x GlutaMax (Thermo Fisher #35050061). Cells were maintained at 37 °C, 5 % O_2_, 10 % CO_2_, and media was replenished every second day. For passaging, 90 % confluent NCCs were dissociated with Accutase (STEMCELL Technologies #07920) for 5-10 minutes and plated at a density of 20,000-30,000 cells/cm^2^, approximately every 3-6 days, until a maximum of 5 passages.

#### EB-NCC Differentiation

Day 0: Aggrewell plates (STEMCELL Technologies #34425, #34415) were prepared using pluronic solution (2% pluronic (Sigma, #P2443) w/v in sterile H_2_O), centrifuging at 1,300 x g for 5 minutes, followed by 2 washes with prewarmed DMEM/F-12. Half differentiation culture volume of NIM + 10 µM ROCKi was added to each well and placed at 37 °C until ready for use. 2 hours before dissociation with Accutase, 10 µM ROCKi was added to hPSCs that contain less than 5% spontaneous differentiation. Cells were incubated in Accutase at 37 °C for 10-20 minutes, until cell aggregates are spontaneously lifting and breaking apart. Accutase was diluted 1:4 in DMEM/F-12 and cell suspension was pelleted at 250 x g for 5 minutes. Cells were resuspended in NIM +10 µM ROCKi and counted on the countess 3 automated cell counter (Thermo Fisher #A49865). Appropriate volumes of cell suspension were added to Aggrewells to generate 500-750cell EBs. This is approximately 3-4.5 x10^6^ cells for the 6 well plate format or 6-9 x10^5^ for the 24well plate format. Wells were then topped up to 6ml (6 well format) or 2.5ml (24 well format) with NIM+ROCKi. Aggrewell plates were centrifuged at 100 x g for 3 minutes and placed at 37 °C, 5 % O_2_, 10 % CO_2_. Day 1:20 to 24 hours after seeding, all media must be carefully removed from each well to minimize disturbing newly formed EBs in microwells. NIM + 10 µM SB431542 (Tocris: 1614) and 500 nM LDN-193189 (STEMCELL Tech. #72147) (NIM+SB/LDN) was carefully added to each well and Aggrewell plates were returned to 37 °C, 5 % O_2_, 10 % CO_2_. Days 2-4: Half media changes with NIM+SB/LDN. Day 5: EBs were dislodged from microwells using a P1000 and transferred to 15 mL conical tubes to allow EBs to gravity settle. Media was refreshed with NIM+SB/LDN, and EBs were plated onto a thin layer (0.16 mg/mL) Matrigel coated TC plastic at a density of 40 EBs / cm^2^. EBs were placed at 37°C, 5% O_2_, 10% CO_2_, dispersed evenly over the growth surface on a leveled shelf. Days 6-11: Media was replaced every other day with NIM+SB/LDN. Day 12: Neutralized adherent EB clusters were carefully washed off the growth surface using a P1000, and media was aspirated. Cells were dissociated with Accutase for 3-5 minutes at 37 °C and dispensed through a 40 µM mesh filter (Falcon #352340) to remove any remaining cell clumps. Accutase was diluted as above, and enriched NCCs were pelleted (250 x g, 5minutes), resuspended in NIM, and seeded onto a thin layer (0.16 mg/mL) Matrigel coated TC plates at a density of 30,000-40,000 cells/cm^2^.

#### NCC Neural/Glial/Mesenchymal Differentiation

For routine passaging, maintenance NCCs were seeded at a density of 20,000-25,000 cells/cm^2^ onto a thin layer (0.16 mg/mL) Matrigel coated TC plates in NIM. Differentiation towards neural, glial, and mesenchymal fates was initiated 24 hours later. Neural: NIM was replaced with basal NIM (bNIM; NIM without insulin, hEGF, and FGF2) supplemented with 50 ng/µL BMP2 (STEMCELL Technologies #78135). Media was fully changed daily for 7days. Glial: NIM was replaced with bNIM supplemented with 20 ng/mL Heregulin-β1 (STEMCELL Technologies #78071) for 5 days. On day 5, media was then changed to bNIM supplemented with 5 µM Forskolin (STEMCELL Technologies #72112) and 10% FBS (Thermo Fisher #12483-020). Mesenchymal: Adipocyte differentiation was carried out using the MesenCult™ Human Adipogenic Differentiation kit (STEMCELL Technologies #05412) according to the manufacturer’s instructions, utilizing the standard mesenchymal stem cell protocol with minimal changes. Briefly, NCCs were plated at maintenance seeding density and left to grow to 90-100% confluence. NIM media was replaced with complete MesenCult™ Adipogenic Differentiation Medium, and cells were differentiated over a period of 14 days with media changes every 2-3 days. Smooth muscle cell (SMC) differentiation was carried out by replacing NIM with bNIM supplemented with 10 ng/mL TGF-β3 (Sigma #SRP3171). After 72 hours, media was replaced with 1:1 blend of bNIM and Media 231 (Thermo Fisher #M-231-500) supplemented with 1 x smooth muscle growth supplement (SMGS (Thermo Fisher #S00725) and 10 ng/mL TGF-β3. After a total of 7 days, media was replaced with Media 231 supplemented with 1X SMGS and 10 ng/mL TGF-β3. Differentiating SMCs were passaged at 90-95% confluence when needed with 0.05% trypsin. Cultures were maintained for an additional 7 days, with media replacement every other day. All cultures were fixed with 3.7% PFA at endpoint for analysis.

#### CRISPR/Cas9 Genome Editing of hPSCs

At 80% confluence, hPSCs were dissociated with Accutase for 10-20 minutes at 37 °C. Cells were rinsed off the growth surface and gently triturated. Accutase was then diluted 1:4 in PBS^—/—^ and then pelleted (250 x g, 5 minutes). hPSCs were resuspended in PBS^—/—^ for counting. For each CRISPR/Cas9 transfection sample, 1 x 10^6^ cells were aliquoted into microcentrifuge tubes and pelleted (250 x g, 5 minutes). Pellets were resuspended in 100µl electrolyte buffer E2 (Thermo Fisher #MPK10096) with 3 µg CRISPR/sgRNA plasmid (Addgene #48139) (Ran et al., 2013) and 1.75 µg of TSC2 exon 3 ssODN donor sequence (see Key Resources Table; Integrated DNA Technologies) or 3 µg *AAVS1:mCherry* donor plasmid (Key Resources Table, supplemental information). Samples were electroporated using the Neon Transfection System (Thermo Fisher MPK5000) using 100 µl tips (Thermo Fisher #MPK10096) at 2 pulses of 1050v, 30 ms pulse width. Electroporated cells were then plated onto diluted Matrigel at approximately 1 x 10^6^ cells into 1 well of a 6-well plate in E8 media containing 10 µM ROCKi. Electroporated cultures were left to grow until individual colonies formed. Expanding colonies were picked into Matrigel coated 96-well cluster plates for PCR/restriction enzyme-based screening for edited clonal cell populations. Karyotypic analysis revealed no chromosomal abnormalities in 7 of the 8 CRISPR/Cas9 edited cell lines utilized in this study (data not shown). Trisomy 12, a common chromosomal abnormality in hPSC cultures (Mayshar et al., 2010), was detected in a subpopulation of H1 *TSC2^—/—^* cells (35%) and is, thus, not related to CRISPR/Cas9 editing. Guide RNA (gRNA) sequences utilized to modify the *TSC2* and *AAVS1* loci (see Key Resources Table) were identified and high scoring (98 and 80, respectively) (Hsu et al., 2013) gRNA sequences were selected to minimize potential off target cleavage events. To identify potential CRISPR/Cas9-induced off-target mutations in *TSC2^—/—^*hPSCs, the highest scoring predicted guide RNA off-target sites were identified using the CRISPOR algorithm (Haeussler et al., 2016). Sequencing of these regions revealed no off-target cleavage induced mutations in any of the *TSC2^—/—^* hPSC lines. See Key Resources Table and supplementary information for DNA/plasmid/oligonucleotide sequences.

#### PCR Screening/Genotyping of CRISPR/Cas9 Edited Cells

Genomic DNA was harvested using QuickExtract solution (screening of CRISPR/Cas9 edited cells; Epicentre #EPI-QE0905T) or NucleoSpin Tissue kit (routine genotyping; Macherey-Nagel #740952.25). OneTaq DNA polymerase kit (New England BioLabs #M0480) was utilized using primers flanking the genome editing target site of exon 3 of *TSC2* (see Key Resources Table for primer sequences). Cycling parameters are as follows: 94°C initial denaturation for 30s, 25x [94°C 20s, 56.5°C 30s, 68°C 30s], 68°C 5m, 4°C ∞. PCR amplicons were then digested with PmeI restriction enzyme (New England BioLabs #R0560S) in OneTaq buffer for 1 hour at 37°C. Agarose gel electrophoresis was utilized to resolve digested amplicons indicating successful integration of ‘stop codon’ donor sequence.

#### Teratoma Assays

Teratoma assays were performed as previously described^92^. Briefly, hPSCs were harvested using Accutase 1 day prior to routine passaging, diluted 1:4 in DMEM/F-12, and pelleted (250 x g, 5 minutes). Cells were resuspended in E8 media and counted. 1 x 10^6^ cells were aliquoted, pelleted, and resuspended in 100 µl ice cold Matrigel for each planned injection site. *NSG* mice were treated with buprenophrine (Richter pharma #02526301) 1 hour before injection, then anesthetized by isofluorane under a continuous stream of O_2_. Matrigel embedded cells were injected bilaterally intramuscularly into the tibialis anterior of NSG mice. Once large visible leg tumors were observed (8-12 weeks), mice were sacrificed and teratomas were excised and fixed in 10% formalin. Teratomas were then processed for paraffin embedding, sectioned, and stained with hematoxylin and eosin for analysis.

#### Imaging Flow Cytometry

hPSCs were dissociated with Accutase, diluted 1:4 in PBS^—/—^, and then pelleted (250 x g, 5 minutes). Cell pellets were resuspended in 3.7% PFA and fixed for 15 minutes at room temperature. Fixed cells were washed twice with PBS^—/—^ in excess, followed by permeabilization in ice-cold 90% methanol for 30 minutes on ice, followed by 2 washes in staining buffer: 1% BSA (Wisent # 800-095-EG), 0.1% Triton-X (BioShop #TRX777) in PBS^—/—^. Samples of 5 x 10^5^ cell were aliquoted into 5ml round bottom tubes and blocked using staining buffer in excess for 30-minutes on ice. Cells were pelleted and resuspended in 100 µl of 1:100 TSC2 antibody (Cell Signaling Technologies, #4308) or 100 µl of 1:100 Cas9 (Cell Signaling, #14697) diluted in staining buffer and incubated at room temperature for 1 hour. Cells were washed in blocking/staining buffer in excess, and resuspended in 100µl of 1:1000 Alexa Fluor 488 Goat Anti-Rabbit IgG (Thermo Fisher, #A11034) or Alexa Fluor 488 Goat Anti-Mouse IgG (Thermo Fisher, #A11001) secondary antibody diluted in staining buffer and incubated for 30 minutes at room temperature, shielded from light. Cells were washed once with staining buffer in excess and resuspended in 200 µl PBS^—/—^ containing 2 µg/ml Hoechst 33342 (Thermo Fisher #H3570). Single stain and no stain controls were prepared in parallel. Cells were passed through a 40 µm cell strainer prior to imaging on the Amnis ImageStreamX Imaging Flow Cytometer.

#### Flow Cytometry of Differentiated NCCs

EB-NCC cultures were rinsed with media to remove neuralized adherent EB-clusters using a P1000, prior to harvesting with Accutase. Matching undifferentiated hPSC lines were harvested in parallel by removing differentiation media containing dislodged EBs, followed by incubation with Accutase for 5 minutes. Cells were gently lifted via trituration, diluted 1:4 in DMEM/F-12, and passed through a 40-µm cell strainer. Cells were pelleted, resuspended in NIM, counted, and 5 x 10^5^ cells were aliquoted into round bottom tubes for staining. Cells were blocked in excess FACs buffer (4% FBS [Thermo Fisher #12483-020], 100 µM EDTA in PBS^—/—^) for 15 minutes on ice. Cells were pelleted and resuspended in 100 µl 1:20 diluted p75-Alexa Fluor 647 conjugated antibody (BD Biosciences: #560326) in FACS buffer. Cells were incubated for 30 minutes on ice, shielded from light. Cells were washed once with excess FACS buffer and resuspended in 500 µl of 1:2000 diluted Sytox Blue nucleic acid stain (Thermo Fisher #S11348) in FACS buffer. Mouse IgG_1_-Alexa Fluor 647 isotype controls (BD Biosciences #557783) and no stain controls were prepared in parallel. Cells were passed through a 40µm filter and immediately measured using the BD Biosciences LSR-Fortessa flow cytometer and analyzed with FloJo (version 10.5.3).

#### Western Blotting

Cell lysates were collected after washing cells with ice cold PBS^—/—^, followed by brief incubation on ice with RIPA cell lysis buffer: 150 mM NaCl (Sigma, #S9888), 1% Triton-X (BioShop, #TRX777), 0.1% SDS (Sigma, #L3771), 50 mM Tris (Sigma, #T1503/#T3253), 1x PhosSTOP phosphatase inhibitor mix (Sigma # 4906845001), 1x complete protease inhibitor mix (Sigma 11873580001). Cells were scraped and transferred to microcentrifuge tubes and rotated at 4°C for 30 minutes, followed by centrifugation at 16,000g at 4°C for 20 minutes to generate cleared lysates. Lysates were stored at -80°C. After thawing for analysis, protein concentrations were normalized and probed via western blot, imaged using the Odyssey Classic Infrared System (LI- COR Biosciences), and analyzed using standard densitometry techniques using FIJI ^1^. Antibodies utilized for western blot are listed in the Key Resources Table.

#### Oil Red O Staining of NCC-derived adipocytes

Oil Red O staining was performed as previously described^93^. Briefly, 2.5 g of Oil Red O (Sigma, #O0625) was thoroughly dissolved in 100% isopropyl alcohol. This stock solution was diluted 1.5:1 in distilled water and then passed through a 45 µm filter. This working solution was then applied to PFA-fixed NCC-derived adipocyte samples for 10 minutes, followed by a 30 minute rinse in tap water. Oil Red O-stained samples were counterstained with hematoxylin for nuclei visualization.

#### Immunofluorescence

Unless specified for specific antibody targets, all immunostaining was performed as follows: Cells were fixed with 3.7% PFA for 20 minutes at room temperature after removing media, followed by 3 washes with PBS^—/—^. Cells were permeabilized in 0.1% Triton-X in PBS^—/—^for 15 minutes and blocked in excess immunofluorescence staining buffer (IF-SB; 1% BSA, 0.1% Tween-20 in PBS^— /—^) for 1h. IF-SB used for blocking was removed, and IF-SB containing diluted primary antibodies (see KRT) was added to each sample and incubated overnight at 4°C. Samples were then washed three times for 5 minutes before the addition of secondary antibody diluted 1:1000 in IF-SB. Samples were incubated in secondary antibody solution for 1 hour at room temperature, followed by three washes with PBS^—/—^, in which 2 µg/ml Hoechst 33342 was included in the final wash step. Antibodies used for immunofluorescence are listed in the Key Resources Table. Images were captured using the Zeiss Observer A1 or Zeiss Observer Z1 fluorescence microscopes or captured for high content analysis using the ArrayScan VTI HCS platform with HCS Studio software or Operetta platform with Harmony software.

Alterations to the immunostaining protocol: HNK-1 cell surface staining was completed after PFA fixation, but prior to cell permeabilization. For PMEL (HMB45) staining, samples were permeabilized using 100% ice-cold methanol for 10 minutes on ice following PFA fixation.

#### O-propargyl-puromycin (OPP) Rate of Protein Synthesis Assay

NCCs were plated at 27,000 cells per cm^2^ onto 24-well plates (Sigma #CLS3473-24EA) and allowed to reach 70-80% confluence. Cells were pulsed with 50 µM O-propargyl-puromycin (OP-puro; MedChemExpress, #HY-15680) for 1 hour. Subsequently, cells were fixed with 4% PFA for 30 minutes. at room temperature. Wells were washed 3x for 5minutes with PBS^—/—^ and permeabilized with 0.1% Triton-X in PBS^—/—^ for 15 minutes at room temperature. The click reaction was prepared by mixing the following components in the described order in PBS^—/—^ to the indicated final concentration: 4 mM CuSO4, 5 µM Sulfo-Cy5-N3 (Luminprobe #A333), 200 mM L-ascorbic acid. The click reaction was mixed, added to the OP-puro treated wells and incubated at room temperature for 30 minutes. Wells were washed 3x for 5 minutes with PBS^—/—^and counterstained with 10 µg/mL Hoechst 33342 for 30 minutes, washed 3x for 5 minutes, and then imaged. Negative controls included untreated cells and cells preincubated with 50 µM cycloheximide (CHX; AdooQ, #A10036) in complete culture medium for 15 minutes. CHX treated media was completely removed, followed by incubation with 50 µM OP-puro. Images were captured for high content analysis using the ArrayScan VTI HCS platform and HCS Studio software.

#### NCC Motility Assay

NCCs were dissociated for plating on day of routine passaging and seeded into 96-well cluster plates at increasing densities of 1,000 to 6,000 cells per well in NIM. 24h later, media was replenished and 100 nM rapamycin or DMSO was added to appropriate samples. Time lapse imaging (15 minute intervals over 6 hours) of mCherry or CellTracker Deep Red (Thermo Fisher #C34565) signal was performed utilizing the Thermo Fisher ArrayScan VTI HCS platform, utilizing the live cell chamber at 37 °C, 5% CO_2_. CellTracker Deep Red staining was performed by staining cells with a working solution of 500 nM in NIM for 30 minutes, followed by 1x wash with PBS^—/—^before replenishing cultures with NIM containing appropriate treatment conditions. Cell tracking image analysis was performed on wells displaying 30-60% confluence, using HCS Studio software.

#### Enzyme-linked immunosorbent assay (ELISA)

Maintenance NCCs at equivalent densities were incubated for 16h in growth-factor deprived media. Conditioned media was collected and centrifuged to remove any cellular debris, then assayed by VEGF-D ELISA kit (R&D Systems, #DY622) with the DuoSet ELISA Ancillary Reagent Kit 2 (R&D Systems, #DY008), following the manufacturer recommended protocols. Briefly, protein-absorbent plates were coated overnight at room temperature using the manufactured-provided capture antibody, followed by 3x washes with PBS^—/—^ + 0.1% Tween-20. We then blocked wells for 1 hour at room temperature with PBS^—/—^ + 1% BSA, then added samples for another 2h incubation at room temperature. After 3x washes with PBS^—/—^ + 0.1% Tween-20, a manufacturer-provided capture antibody was added for 2h at room temperature, followed by 3x washes. Finally, wells were incubated with the provided streptavidin-HRP conjugate for 20 minutes, washed 3x, and subjected to a colorimetric reaction with H_2_O_2_ and tetramethylbenzidine. After 20 minutes, the colorimetric reaction was stopped with 2N H_2_SO_4_, and the optical density at 450 nm was determined for each well. Calculations for sample VEGF-D concentrations were determined by comparing against a standard curve.

#### RNA-Seq sample preparation and sequencing

Cells were dissociated using Accutase and diluted 1:4 in PBS^—/—^ prior to pelleting (250 g, 5 minutes). Cells were resuspended in PBS^—/—^ and total cell count was noted. Cells were pelleted (250g, 5minutes) and resuspended in TRIzol reagent (Thermo Fisher # 15596026). Samples were stored immediately at -80°C until processing for RNA extraction. RNA extraction was performed in batches which included all differentiation time points within a WT and *TSC2^—/—^* cell line pairing. RNA extraction was performed according to the manufacturer’s protocol followed by DNase digestion (Qiagen #79254) and ethanol precipitation of RNA. RNA concentration was estimated using NanoDrop 2000 spectrophotometer (Thermo Fisher ND-2000) and ERCC spike in (Thermo Fisher #4456740) was added, normalizing to cell number. Quality control of RNA integrity and total RNA concentration was then measured using the Bioanalyzer Eukaryote Total RNA Pico kit (Agilent #5067-1513). Only samples with RIN values of 8.0 or greater were submitted for sequencing. Total RNA H9 WT and *TSC2^—/—^* NCC sample libraries were prepared with ribosome depletion (Illumina # 20020598), all other sample libraries were prepared using the TruSeq Stranded mRNA kit (Illumina #RS-122-2101). All samples were sequenced on the Illumina NextSeq500 or NovaSeq6000 platform (Illumina) generating at least 30 million reads per sample. Reads were assigned to transcripts from GENCODE release 39 using Salmon (Patro et al., 2017). All RNA sequencing datasets are available on GEO (accession #GSE137614).

#### Differentially expressed genes (Time Course)

Differentially expressed genes were identified using DESeq2 v1.42.1 (Love et al., 2014). DESeq2 analysis modelled fold change between time points using the cell line as an independent factor (using the model ’*∼cell_line + condition*’), and fold changes were calculated using lfcShrink function, applying the apeglm method (v 1.2.1) (Zhu et al., 2018). Gene ontology enrichment analyses of differentially expressed genes were performed using clusterProfiler R package (Yu et al., 2012).

#### Differentially expressed genes (Sex-dependent differences)

Pseudocount abundance data generated by *Salmon* was imported into the *DEseq2* framework in R for differential expression and enrichment analysis. Principal components analysis was conducted on all samples to visualize transcriptomes in a two-dimensional space. For single variable differential expression testing, we subsetted samples to only include the untreated and fit the following model: *∼ batch + genotype + sex*. We then tested for genes with significant coefficients by Wald test, separately for genotype and for sex. To assess for changes across genotype that differs between sex, we again subsetted for untreated samples and fit the following model: *∼ batch + genotype + sex +genotype:sex*. The interaction term coefficient for each gene was tested for significance by Wald test. Differentially expressed genes were called when false discovery rate (FDR) < 0.05, ± |log_2_FoldChange| > 1 (as indicated in the text). To visualize expression values by heatmap, sample conditions were collapsed by abundance summation, normalized, and then transformed by regularized log_2_ transformation (implemented in *DEseq2*). Heatmaps were generated using the *pheatmap* package in R. KEGG pathway enrichment was performed using *g.Profiler*^94^ on significant DEGs (FDR < 0.05, ± |log2FoldChange| > 1) and visualized using Cytoscape. All analysis was conducted using R 4.0.3 within the RStudio 023.12.1environment.

#### Differentially expressed genes (ASO treatments)

Pseudocount abundance data generated by *Salmon* was imported into the *DEseq2* framework in R for differential expression and enrichment analysis. Principal components analysis was conducted on all samples to visualize transcriptomes in a two-dimensional space. For single variable differential expression testing, we subsetted samples to only include the untreated and fit the following model: *∼ batch + genotype + treatment*. We then tested for genes with significant coefficients by Wald test, separately for genotype and for treatment. To assess for changes across genotype that differ between treatment, we again subsetted for untreated samples and fit the following model: *∼ batch + genotype + sex +genotype:treatmeant*. The interaction term coefficient for each gene was tested for significance by Wald test. Differentially expressed genes were called when false discovery rate (FDR) < 0.05, ± |log_2_FoldChange| > 1 (as indicated in the text). To visualize expression values by heatmap, sample conditions were collapsed by abundance summation, normalized, and then transformed by regularized log_2_ transformation (implemented in *DEseq2*). Heatmaps were generated using the *pheatmap* package in R. GO term enrichment was performed using *g.Profiler*^94^ on significant DEGs (FDR < 0.05, ± |log2FoldChange| > 1) and visualized using Cytoscape. All analysis was conducted using R 4.0.3 within the RStudio 023.12.1environment.

### Genome wide synthetic lethal screen

#### Generation of Cas9-expressing WT and TSC2^—/—^ NCCs

To generate Cas9 expressing cell linces, WT and *TSC2^—/—^* H9 hPSCs were transduced with the lentiCas9-Blast lentiviral vector ^95^. Transduced cells then underwent blasticidin selection to enrich for transduced cells before subcloning to establish uniform populations of Cas9 expressing cells.

#### GeCKOv2 Lentiviral Library Preparation

The human GeCKOv2 libraries^96^ A & B (Addgene #1000000049) were diluted to 5 ng/µl in sterile ddH2O for electroporation. 2 µl of Library A or B were added to 25 µl of 5-alpha electrocompetent cells (New England Biolabs #C2989K) were added to a prechilled 1mm electroporation cuvette, and cells were electroporated (4 per GeCKOv2 library) using the GenePulser electroporator (Bio-Rad) using the following parameters: 1.7 kV, 200 Omega, and 25 μF. 1ml of SOC outgrowth media (New England Biolabs #B9020S) was added immediately following electroporation, and cells were transferred to 17 mm x 100 mm round-bottom culture tubes for recovery in a bacterial shaker 250 rpm at 37°C for 1h. Each respective library was then combined. 10µl of recovered bacteria from each library was diluted into 1 ml of SOC. 20 µl of or diluted cells were spread onto pre-warmed ampicillin LB-agar plates. Plates were incubated overnight at 32°C. 1ml of LB was added to each plate and bacterial colonies were scraped, combined, and pelleted at 6,000 g for 5 minutes, and then weighed. One maxiprep (Thermo Fisher # K210016) column per 500 mg bacteria was utilized to isolate plasmid DNA.

#### Lentivirus production and determination of functional titer

HEK 293T cells were seeded in 150 mm^2^ tissue culture dishes (Corning # 353025) for transfection. Cells were cotransfected with second-generation lentiviral packaging plasmids pMD2G, pPAX2, and the GeCKOv2 libraries A&B, lentiCas9-Blast (Addgene67 #52962), or lentiGuide-Puro (Addgene #52963) in the presence of polyethyleneimine (4.1 μmol/L) and NaCl (2.25 × 10−4 mol/L). GeCKOv2 and lentiCas9-Blast virus was produced using 40x15-cm dishes for each virus construct, and lentiGuide-Puro virus with 1x15 cm^2^ dish. Supernatant containing the virus was collected at 48h after transfection and immediately aliquoted and stored at -80°C. Functional titers were established as follows: Virus supernatant was thawed in warm rapidly in warm water. *WT* or *TSC2^—/—^* NCCs were plated into 6x PLO/Fn coated 100 mm^2^ dishes at 1.2 x 10^6^ cells per dish including 500, 250, 125, or 62.5 µl of virus supernatant, leaving 1 dish as selection control and 1 dish as a no treatment control. 24 hours post transduction, transduced cells and one not treatment control were administered selection (1 µg/ml puromycin (Thermo Fisher # A1113803) or 5µg/ml blasticidin (InvivoGen #ant-bl-1)) for 48 hours. Cells were then harvested and counted, comparing no treatment control to transduced samples to establish the number of functional virus particles per µl of virus supernatant.

#### Synthetic Lethal Screen

NCCs were differentiated on a massive scale in the same manner as described above. For optimal GeCKOv2 transduction at 700X representation, 2.9 x 108 NCCs were combined with enough virus supernatant to allow 8.7 x 10^7^ NCCs to be transduced (30% of seeded) and seeded into Matrigel coated 150 mm^2^ dishes (7.595 x 10^6^ NCCs in 16.5 ml NIM media per dish). 2x 100 mm^2^ dishes were seeded in addition to determine experimental MOI for representation calculations (# cells/123,411 gRNAs). 24 hours post transduction, media was changed to NIM with 1 µg/ml puromycin (1x 100mm^2^ dish was not selected with puromycin) and NCCs were selected for 72 hours. NCCs within the 100 mm^2^ dishes were counted to determine transduction efficiency (30-50% ideal, at least 700X representation). For d0 baseline time points at least 61.7 x 10^6^ NCCs were collected and pelleted at 500g for 5 minutes and then placed at -80°C for storage. Media was replenished in remaining cultures, which were maintained per usual in 150 mm^2^ dishes, ensuring at least 1000X representation for 14 days. At endpoint, cells were harvested using Accutase, counted, and pelleted at 500g for 5 minutes, and stored at -80°C for downstream processing for deep sequencing of genomic integrations.

#### Genomic DNA isolation: Synthetic Lethal Screen

Frozen samples were divided into 30-50 x10^6^ per 15 ml centrifuge tubes. For each tube, 6 ml of NK Lysis Buffer (50 mM Tris, 50 mM EDTA, 1% SDS, pH 8) and 30 μl of 20 mg/ml Proteinase K (Qiagen #19131) were added to the cell sample and incubated at 55 °C overnight in a water bath. The following day, 30 μl of 10 mg/ml RNAse A (Qiagen #19101, diluted in NK Lysis Buffer to 10 mg/ml) was added to the lysed sample. Tubes were then inverted at least 25 times and incubated at 37 °C for 30 minutes. Samples were then immediately cooled on ice before addition of 2 ml of pre-chilled 7.5 M ammonium acetate (Sigma #A1542) to precipitate proteins. Stock solutions of 7.5 M ammonium acetate were made in sterile ddH_2_O and kept at 4 °C until use. Samples were then vortexed at high speed for 20 seconds and followed by centrifugation at ≥ 4,000 x g for 10 minutes. Supernatant was then carefully decanted into a new 15ml centrifuge tube. 6 ml of 100% isopropanol was then added and inverted at least 50 times before centrifugation at ≥4,000 x g for 10 minutes. The supernatant was then discarded and pelleted DNA was washed twice with 70% ethanol (≥4,000 x g for 1 minute for each wash). Ethanol was carefully removed, and DNA pellets were air dried for 10-20 min. DNA was resuspended in 1x TE buffer (Sigma #9285) incubated at 65°C for 1h and left at room temperature overnight to fully dissolve. The gDNA concentration was measured using the Nanodrop Micro volume Spectrophotometer (Thermo Scientific).

#### Library Preparation and Sequencing Parameters: Synthetic Lethal Screen

PCR1: To maintain proper representation for deep sequencing of lentiviral integrations, 450 µg of gDNA per time point was used as a template and split into 50µl reactions using the NEBNext® Ultra II Q5 Master Mix (New England Biolabs #M0544), using the GeCKOv2 Adaptor_F and GecKOv2 Adaptor_R primers, using the manufacturers recommended reaction composition and thermocycling parameters. Cycling parameters were as follows: 98°C initial denaturation for 30s, 13x [98°C 10s, 65°C 75s,], 65°C 5m, 4°C ∞. For each time point, PCR reactions were combined into a single 15ml centrifuge tube and mixed well, to serve as a template for PCR2 reactions to generate Illumina adaptors to PCR1 amplicons. PCR2: For each time point, 13x 50 µl reactions of NEBNext® Ultra II Q5 Master Mix were prepared using 2 µl/50 µl reaction of PCR1. All time points used the reactions used the GeCKO_Master_F forward primer combined with unique reverse primers for each sample to barcode each time point separately for multiplexing. The thermocycling parameters were identical to PCR1. PCR2 reactions from each time point were pooled and then gel purified to attain the desired ∼360 base pair amplicons using the QIAquick Gel Extraction Kit (Qiagen #28704) following the manufacturer’s protocol except gels were dissolved at 30°C. Library concentrations were then estimated using the Nanodrop (Thermo Scientific). Samples were stored at -20°C until further processed. Sample were sequenced with 15% Phi-X spike in on the NextSeq500 platform (Illumina) using High Output chemistry using 30 dark cycles, followed by 20 imaging cycles, and 8 index read cycles. A minimum 60 million reads for day 0 samples, and minimum 25 million reads for day 14 samples were sequenced. Data analysis was carried out using MAGeCK-VISPR , MAGeCKFlute and BAGEL software packages^97–100^. GO term and KEGG pathway enrichment analysis was performed with g.Profiler^94^.

### ASO assays

#### PCR for ASO knockdown validation

Culture media was aspirated from wells and cells were immediately lysed with TriZOL. 200 μL of chloroform was added followed by centrifugation at 15,000g for 20 minutes at 4°C. The colourless aqueous phase was extracted and mixed with equal volumes of 70% EtOH. Following a brief incubation at room temperature, the solution was eluted through a Macherey-Nagel Nucleospin column. We proceeded with column-based purification as per manufacturing protocol. Samples quality was assessed by Nanodrop, and only samples with A260/A280 of 1.8-2.3 and A260/230 > 2 were used. We reverse-transcribed 1µg of total RNA from each sample via SuperScript II with oligoDT priming following manufacturer instructions. qPCR was performed using SYBR Green I and analysis on LightCycler® 480 Instrument II (Roche). We quantified normalized gene expression by scaling Cq by primer efficiency, dividing by the geometric mean of reference gene values (ACTB and GAPDH), and normalizing by the within-group average value.

#### Cell treatments

All ASOs were resuspended in TE buffer: 10 mM Tris, pH 7.5, 0.1 mM EDTA, and diluted in NIM before adding to wells with cells. Across all experiments what were done in the presence and absence of rapamycin, it was consistently used at a 100nM concentration and added 16hours after treatment with ASOs. All compound treatments were conducted for 72 hours unless otherwise stated.

ASO treatments were conducted via gymnosis for 72 hours, unless otherwise stated. 16 hours after ASO treatment 100nM of rapamycin was added. Live cell imaging dyes (Hoechst, SyTOX and Alamar Blue) were incubated for 30 minutes. Prior to imaging, dyes were added as 3X concentrates in NIM to avoid cell detachment. Live imaging dyes were not washed prior to imaging; this did not impact image acquisition as dyes are minimally fluorescent unless bound to the target molecule.

#### EdU proliferation assay

Cells were pulsed with 5 µM of EdU for 3 hours. Subsequently, cells were fixed with 4%PFA for 20 minutues at room temperature. Wells were washed 2 x15 minutes with PBS, permeabilized with 0.1% Triton-X in PBS for 20 minutes at room temperature, then washed again 3 x 5 minutes. Click reaction was prepared by mixing the following components in the described order, in PBS, to the indicated final concentrations: 4 mM Cu2SO4, 5 µM Sulfo Cy5-N3, and 100 mM L-ascorbic acid. The click reaction mix was added to wells containing cells and incubated at room temperature for 30 minutes. Wells were washed 1 x 20 minutes with PBS, counterstained with 10 µg/mL Hoechst 33342 for 30 minutes, washed again 1 x 20 minutes, and then imaged.

#### Clonogenic assay

Cells were plated on Matrigel and treated with ASO at 500 nM via gymnosis every 5 days. After 10 days, cells were dissociated with Accutase for 3-5 minutes then plated on Matrigel, in serial dilution. Cells were permitted to proliferate for 10-15 days, forming colonies from single cells. Following, wells were fixed with 4% PFA for 15 minutes at room temperature, then washed 3 x 5 minutes with PBS. Colonies were stained with 0.1% crystal violet for 1 hr at room temperature, washed 3 x 5 minutes with ddH_2_O, air dried, and imaged.

#### Micronuclei

Micronuclei were quantified by high content imaging-based analysis using the operetta imaging system and harmony software of fixed cultures. For each well, the total number of micronuclei-positive events was counted and normalized to the total number of intact nuclei within the same field of view to account for differences in cell density. Intact nuclei were defined based on size and morphology consistent with non-fragmented, Hoechst-positive nuclear structures. The micronuclei number was calculated as the ratio of micronuclei to total nuclei per well. Values were subsequently normalized to the mean micronuclei of scramble ASO control samples to enable comparison across conditions.

#### 3D NCC spheroid invasion assay

The protocol were performed as previously described^101^ with modifications . Briefly, cells were seeded into Corning 96-well spheroid microplate (Corning, #4520) at 1000cells per well in 100 μL of NIM and incubated 24h to produce spheroids. The following day they were imaged using high content imaging (PerkinElmer Operetta CLS) to obtain initial size of spheroid in each well prior to being treated with ASO and overlaid with Matrigel. After acquiring images spheroids are treated with 20 μL of ASO in NIM at 12X the final desired concentration and immediately after 120 μL of Matrigel at 2X the final desired concentration is overlaid into the well. Cells were allowed to invade for 48-72 hours and were imaged using PerkinElmer Operetta CLS. Invasion was quantified in ImageJ by measuring the area of invading cells relative to the area of the spheroid core.

#### 105k Allograft

Cells were harvested using 0.05% trypsin, resuspended, and diluted to a final concentration of 1.5 × 10⁴ cells/µL in a 1:1 mixture of serum-free DMEM and Matrigel. The cell suspension was maintained on ice (4°C) until injection. For tumor implantation, 1.5 × 10⁶ cells (100 µL per injection) were administered subcutaneously into the left and right hind flanks of *C57BL/6* mice (n = 25). Prior to injection, mice were anesthetized with isoflurane using an induction chamber and maintained via nose cone. The injection sites were shaved and sterilized with isopropyl alcohol. Injections were performed using 1 mL syringes fitted with 25-gauge needles. Following implantation, mice were removed from anesthesia and allowed to recover on a heating pad before being returned to their home cages. Mouse weights and caliper measurements were taken daily. Tumor volume was calculated as 0.5xLengthxWidth^2^. Drug treatment began 11 days post implantation, when all tumors had reached 100mm^3^, and continued over a 10-day treatment period or 30-day treatment period, according to the dosing regimen in figure legends. At experimental endpoint all mice were euthanized, and the tumors were extracted, weighed, and volumes measured by H_2_O displacement.

#### Use of publicly available datasets for gene signature analysis

The gene signature of Pulmonary LAM cells and mouse cranial and trunk NCCs was utilized as presented (Guo et al., 2019; Soldatov et al., 2019). All other gene signatures were established as the significantly up-regulated genes presented within each publication: TSC-associated tumors rAMLs, 621-101 cells^30^ & *Tsc2^—/—^* MEFs^31^, *Tsc2^—/—^* NDFs^32^.Mouse gene signatures were converted to human orthologs prior to analysis. Gene set overlap analysis was performed using the GeneOverlap R package and comparative analysis was performed using the clusterProfiler R package.

#### Statistical analysis

All figures are presented with individual data points (where graphically appropriate), with measures of central tendency and error to be mean and standard deviation, respectively, unless otherwise stated. Sample sizes (n), statistical testing procedures, and post-hoc analyses employed are also reported in each figure legend, with significance (*) attributed when p < 0.05. In general, for normally distributed data sets with equal variances, we performed unpaired two-tailed t-tests, or one-way or two-way ANOVA testing followed by Bonferroni or Tukey’s post-hoc testing. When data were normalized, it was done to their respective control. Sample sizes for both *in vitro* and *in vivo* studies were determined according to field-specific conventions. For every experiment, we had at minimum an n = 2 for each genotype (two independent replicates from H9, H7, H1 and D168). Experiments were repeated three or more times unless otherwise noted, with replicates collected under independent conditions. Outliers were only excluded if there was definitive empirical evidence of technical error. Software for statistical analyses include GraphPad Prism 11 and the programming language R v4.3.2.

---

**Figure S1:**
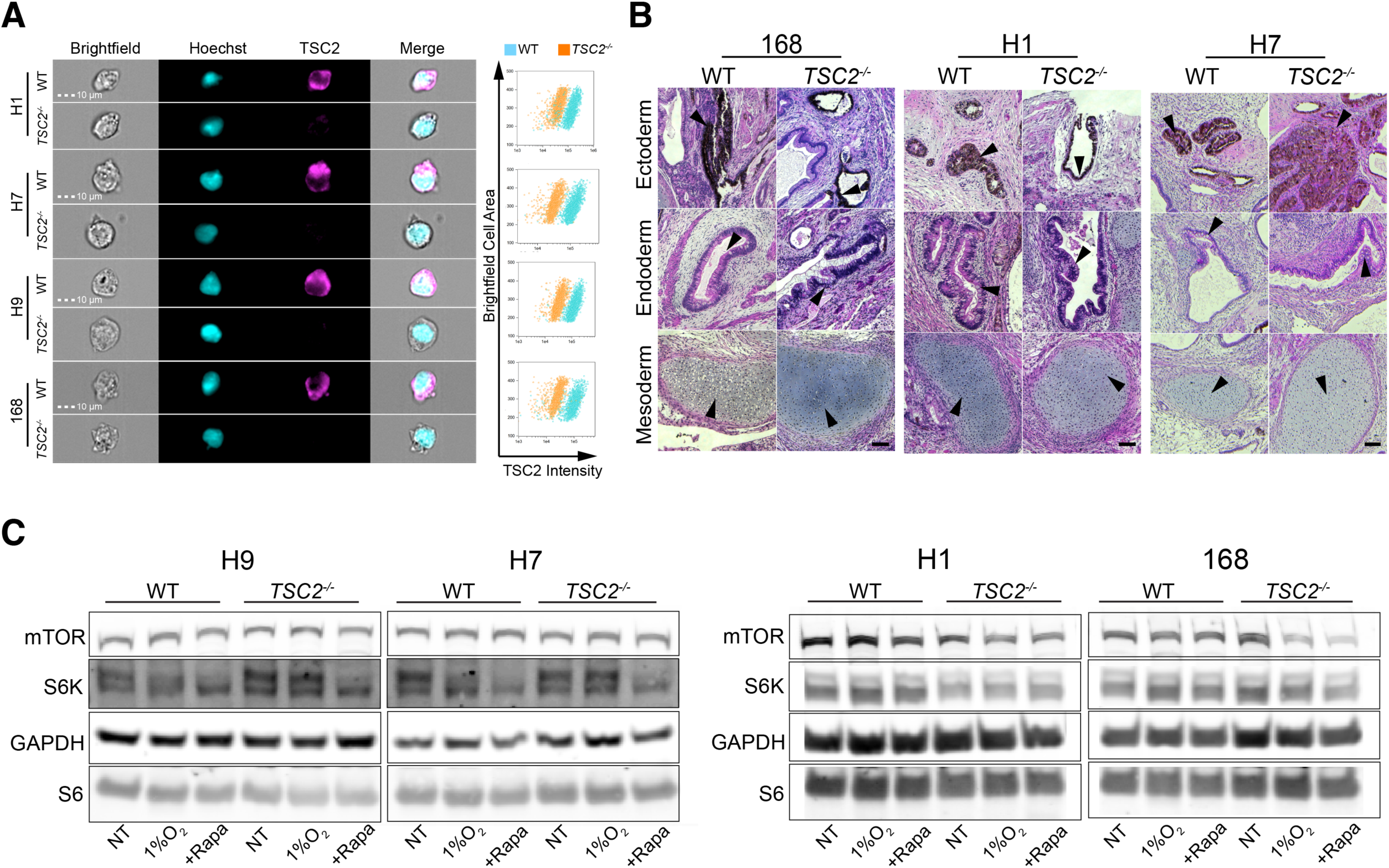
Validation of TSC2 knockout in hPSCs. (A) Imaging flow cytometry image capture following staining with TSC2 antibody confirm clonal populations of TSC2-null cells. (B) H&E staining of 168, H1 and H7 WT and *TSC2^—/—^* teratomas. Arrows indicte examples of tissues of ectodermal, endodermal, and mesodermal origin. Scale bar, 100 µm. (C) Western blots probing mTOR, S6K, and S6 of samples treated for 6h under no treatment, 1% O2, and 100 nM rapamycin conditions.

**Figure S2:**
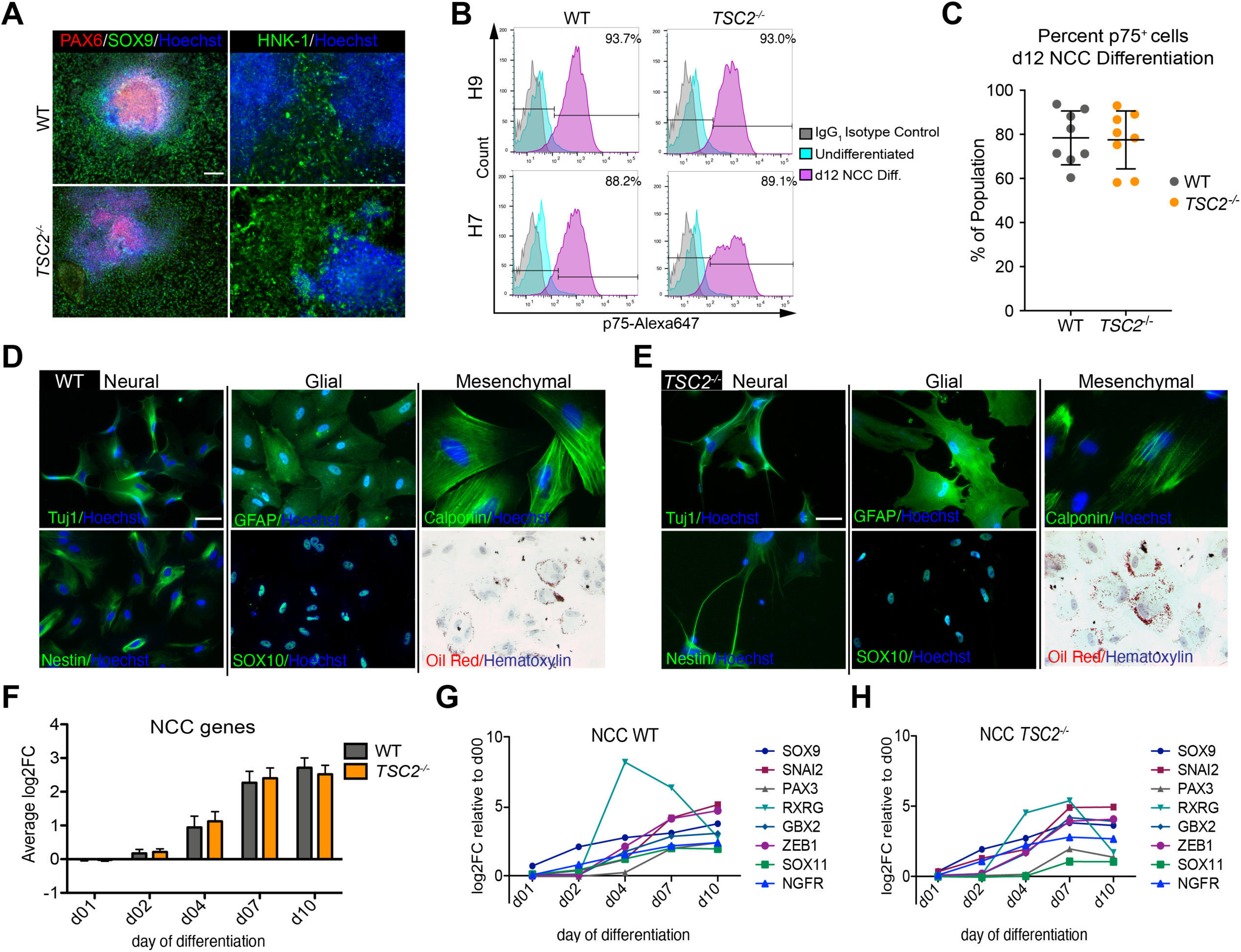
*TSC2^—/—^*and WT EB-NCC differentiation yields enriched and multipotent NCCs. (A) Representative immunofluorescence staining of lineage specific markers PAX6 (neural ectoderm), SOX9 and HNK-1 (NCC) in WT and *TSC*/— cultures at day 12 of the EB-NCC differentiation. Scale bar, 100 µm. (B) Representative flow cytometry analysis of p75 expression in WT and *TSC2*^—^/— cultures at day 12 of the EB-NCC differentiation prior to establishing maintenance populations. (C) Percent p75^+^ cells at day 12 of the EB-NCC differentiation PAX6 (mean ± SD; n = 8 [n = 2 of 168, n = 2 of H9, n = 2 of H7, n = 2 of H1]). No statistical significance was observed between WT and *TSC2^—/—^*NCCs. (D&E) Representative immunofluorescence staining of WT and *TSC2^—/—^*NCCs differentiated towards neural (neuronal markers: Tuj1, Nestin), glial (Schwann cells: GFAP, maintaining SOX10 expression at endpoint) and mesenchymal lineages (smooth muscle cells: Calponin; adipocytes: oil red staining). Scale bars, 50 µm. (F) Average Log2FC of eight neural crest lineage genes (SOX9, SNAI2, PAX3, RXRG, GBX2, ZEB1, SOX11, NGFR) throughout NCC differentiation. Values represent mean Log2FC ±SEM of all eight genes at each time point relative to WT at day 0. (G-H) Expression of individual neural crest lineage genes throughout NCC differentiation with (G) representing WT NCCs and (H) representing *TSC^—/—^* NCCs.

**Figure S3:**
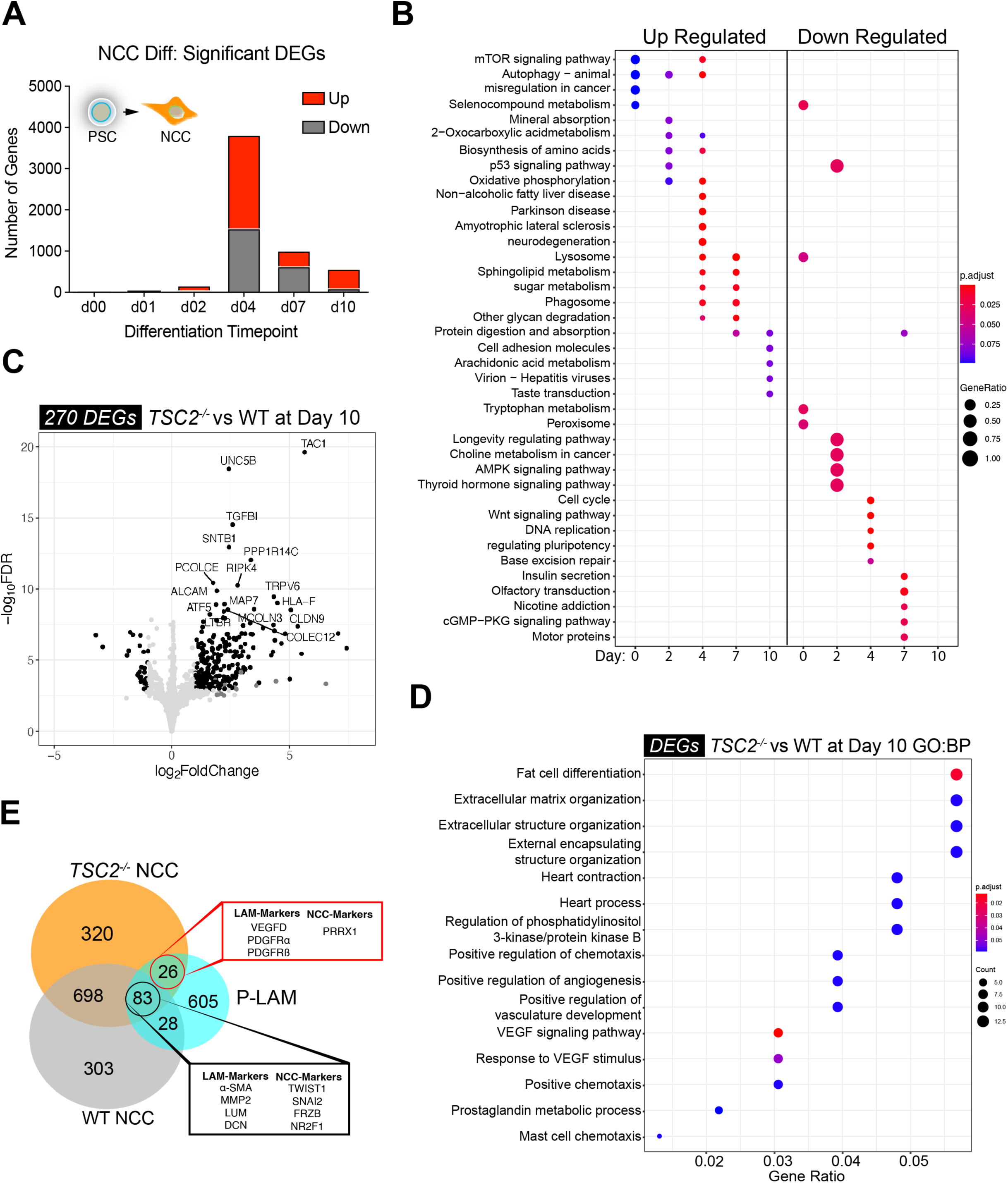
RNA-seq analysis of day 10 *TSC2^—/—^* NCCs exhibit transcriptional features of LAM. (A) Number of total DEGs identified (FDR < 0.05 and |log2FC| > 1) over the RNA-seq differentiation time-course. (B) Dotplot of KEGG enrichment analysis of significantly upregulated and downregulated DEG lists (FDR < 0.05 and |log2FC| > 1) throughout the RNA-seq differentiation time-course. (C) Volcano plot upon comparing *TSC2^—/—^* vs. WT NCC at day 10 of EB-NCC differentiation. Black points are considered differentially expressed (FDR < 0.05 and |log2FC| > 1). The 20 most significantly expressed DEGs are noted. (D) Dotplot of GO term enrichment analysis of DEG lists (FDR < 0.05 and |log2FC| > 1) upon comparing *TSC2^—/—^* vs. WT NCC at day 10 of EB-NCC differentiation. Top 15 most significantly enriched terms are plotted. (E) Venn diagram featuring the intersection of the protein coding gene signatures of WT NCCs, *TSC2*^—/—^ NCCs, and LAM cells.

**Figure S4:**
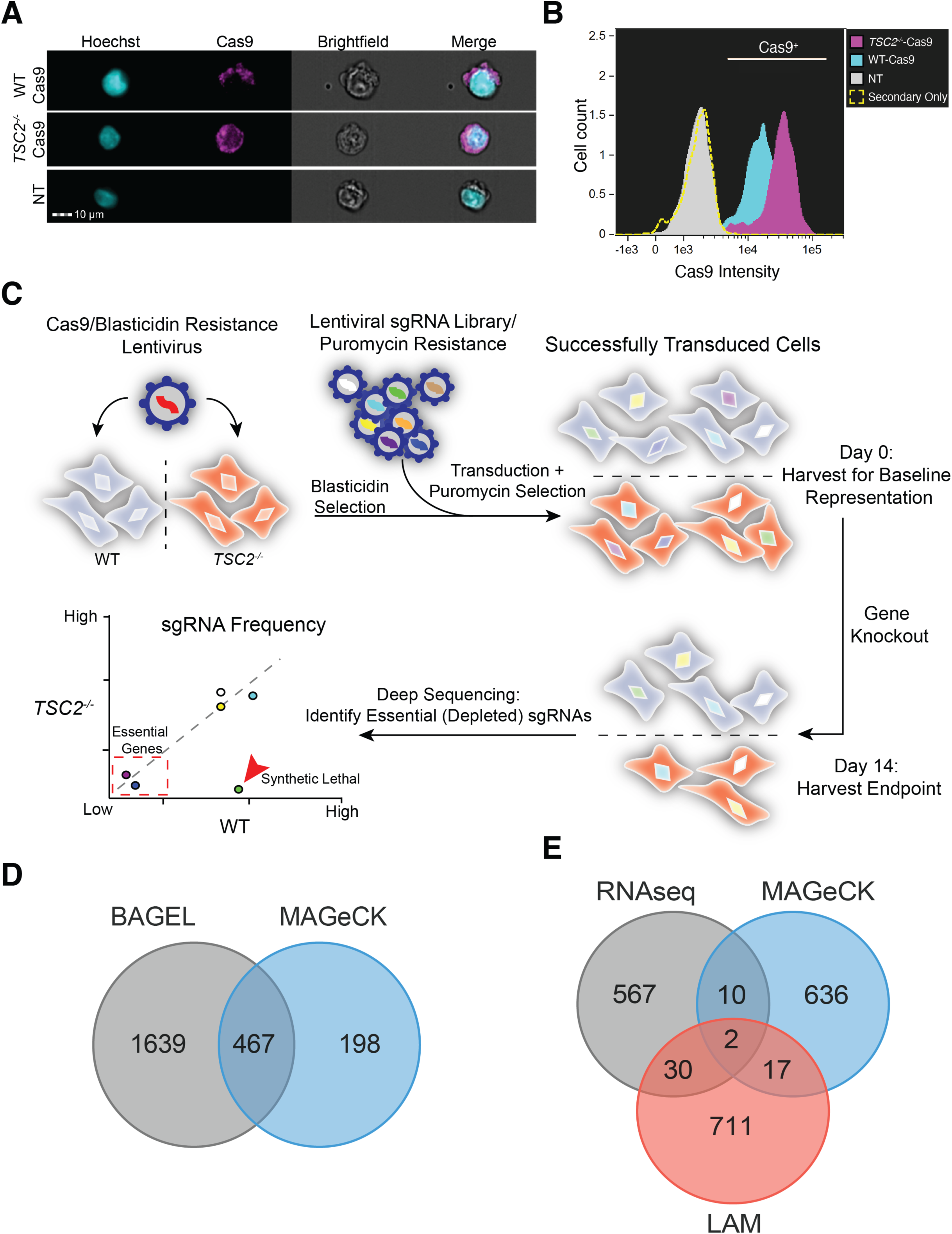
Functional validation of the GeCKO assay system. (A) Imaging flow cytometry image capture of Cas9 expressing H9 WT and *TSC2^—/—^* cell lines. Scale bar, 10 µm. (B) Histogram of imaging flow cytometry analysis displaying enriched population of Cas9 expressing cells in H9 WT and *TSC2^—/—^*cell lines. (C) Schematic representation of experimental workflow of the dropout GeCKOv2 CRISPR/Cas9 screen in WT and *TSC2^—/—^* NCC. (D) Venn diagram of the overlap of top synthetic lethal genes Identified by two different algorithms, BAGEL and MAGeK. (E) Overlap of DEGs identified from day 10 *TSC2^—/—^*vs. WT NCCs, LAM genes, and group IV synthetic lethal genes.

**Figure S5:**
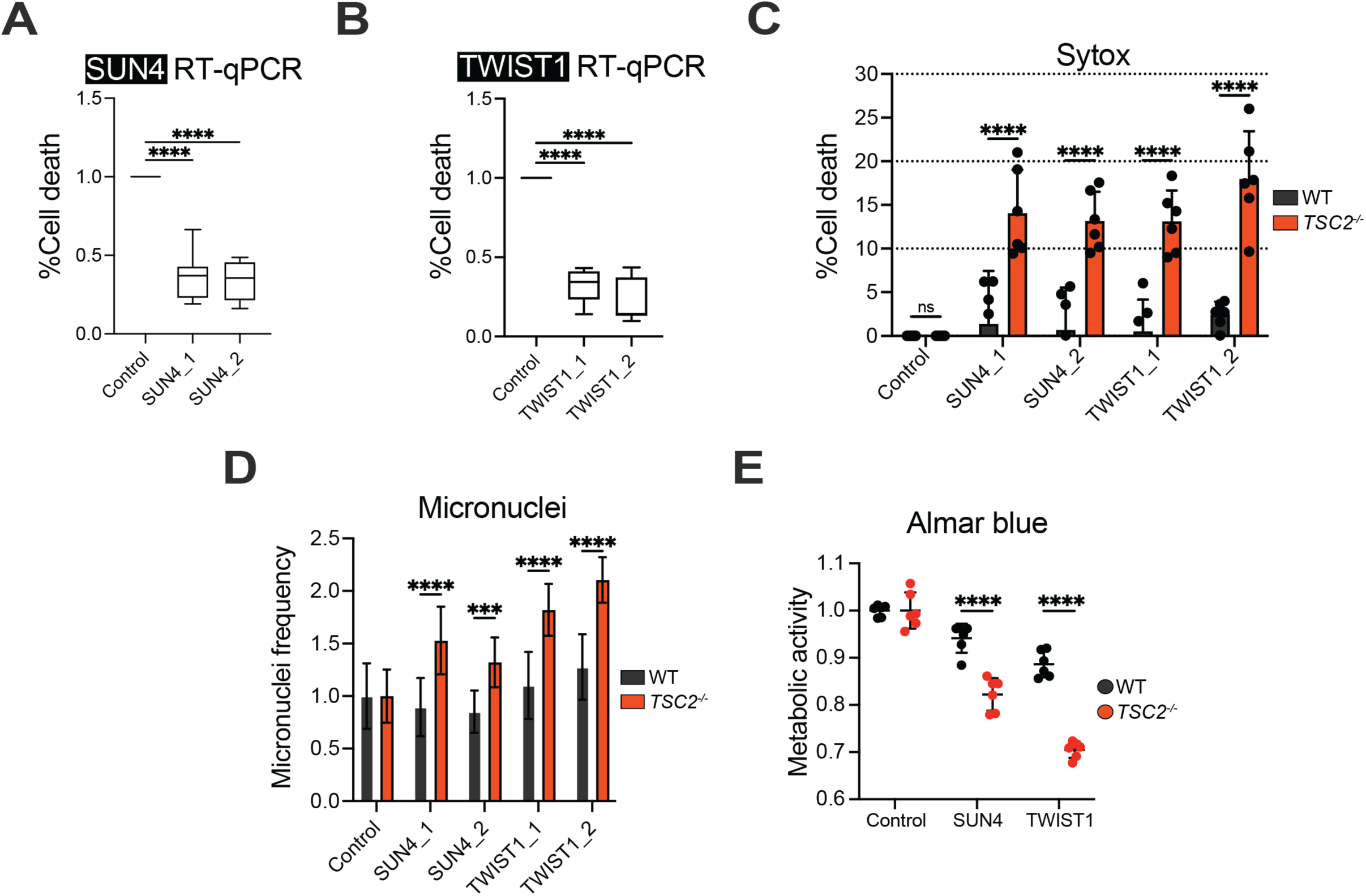
TWIST1 and SUN4 inhibition induce synthetic lethality in *TSC2^—/—^* NCCs. (A&B) Validation of knockdown via quantitative RT-PCR analysis of TWIST1 and SUN4 following 48 hours of ASO treatment at 500 nM. (mean ± SD; n= 6 [n = 3 of H9, n = 3 of H7]; ∗p < 0.05 by using one-way ANOVA and Tukey’s post hoc analysis; Values are normalized to Control-ASO). (A) SUN4-ASO (SUN4_1, SUN4_2). (B)TWIST1-ASO (TWIST1_1, TWIST1_2). (C) Percentage of Sytox^+^ cells after 72 hours treatment with ASOs (mean ± SD; n= 6 [n = 3 of H9, n = 3 of H7]; ∗p < 0.05 by using two-way ANOVA and Tukey’s post hoc analysis; Values are normalized to Control-ASO within genotype). (D) Micronuclei frequency in NCC treated with Control, TWIST1 or SUN4 ASOs for 48 hours (mean ± SD; n= 6 [n = 3 of H9, n = 3 of H7]; ∗p < 0.05 by using two-way ANOVA and Tukey’s post hoc analysis; Values are normalized to Control-ASO within genotype). (E) Cell metabolic activity assessed with Alamar blue 48 hours after ASO treatment (mean ± SD; n= 6 [n = 3 of H9, n = 3 of H7]; ∗p < 0.05 by using two-way ANOVA and Tukey’s post hoc analysis; Values are normalized to Control-ASO within genotype).

**Figure S6:**
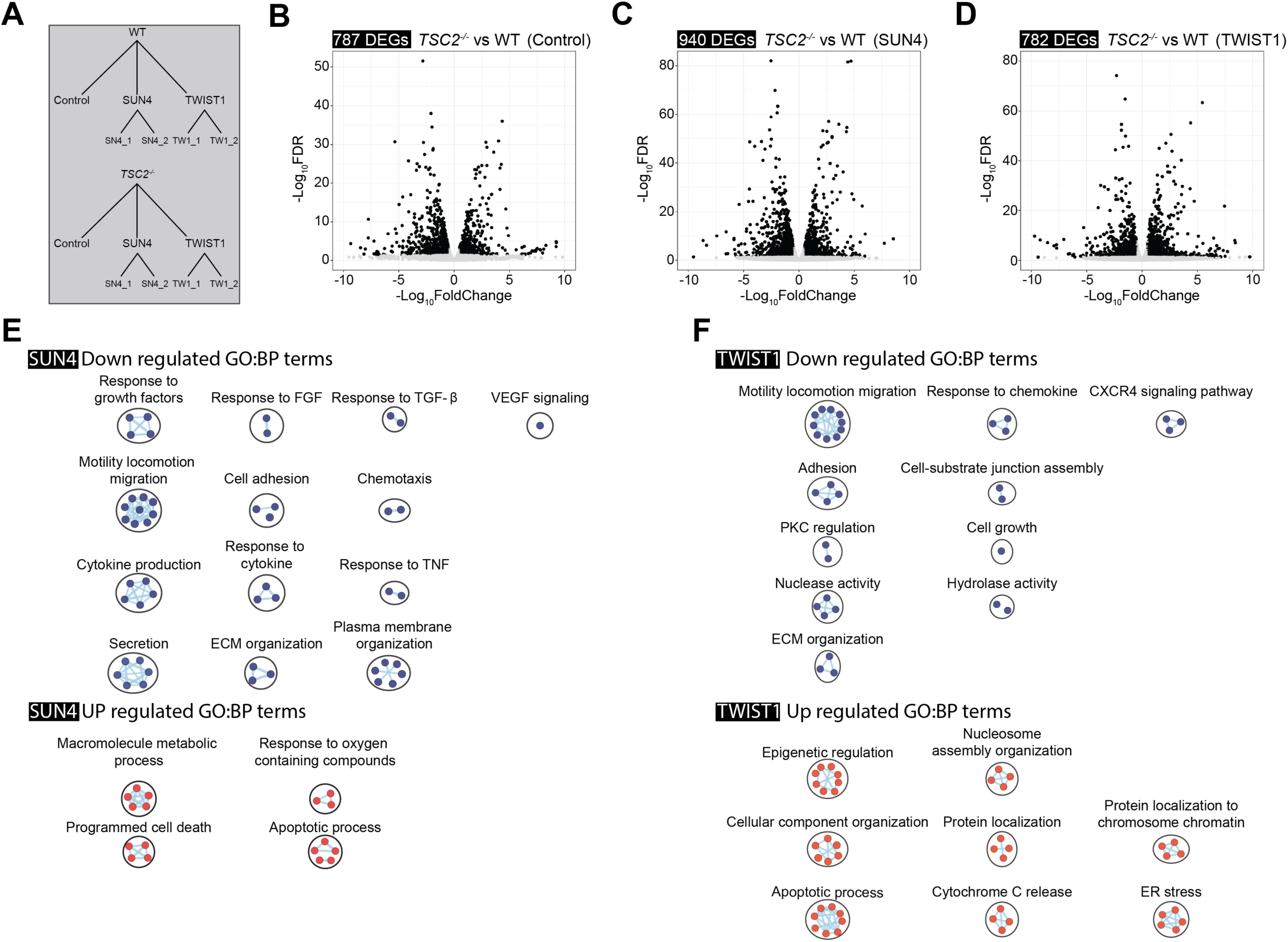
TWIST1 and SUN4 reverse metastatic effects of *TSC2^—/—^* cells. (A) Schematic of the sample conditions tested in the bulk RNA-seq experiment. All cells were treated for 48 hours with 500 nM of ASO (B-D) Volcano plot comparing *TSC2^—/—^* vs. WT cells for Control-ASO treated (B), (C) TWIST1 treated and (D) SUN4 treated cells. Points highlighted in black are considered differentially expressed (FDR < 0.05 and |log2FC| > 1). (E&F) GO term enrichment analysis was performed on genes showing treatment-dependent expression changes to *TSC2^—/—^* condition (FDR < 0.05 and |log₂FC| > 1). Enriched terms were clustered in Cystoscape based on shared biological process.

**Figure S7.**
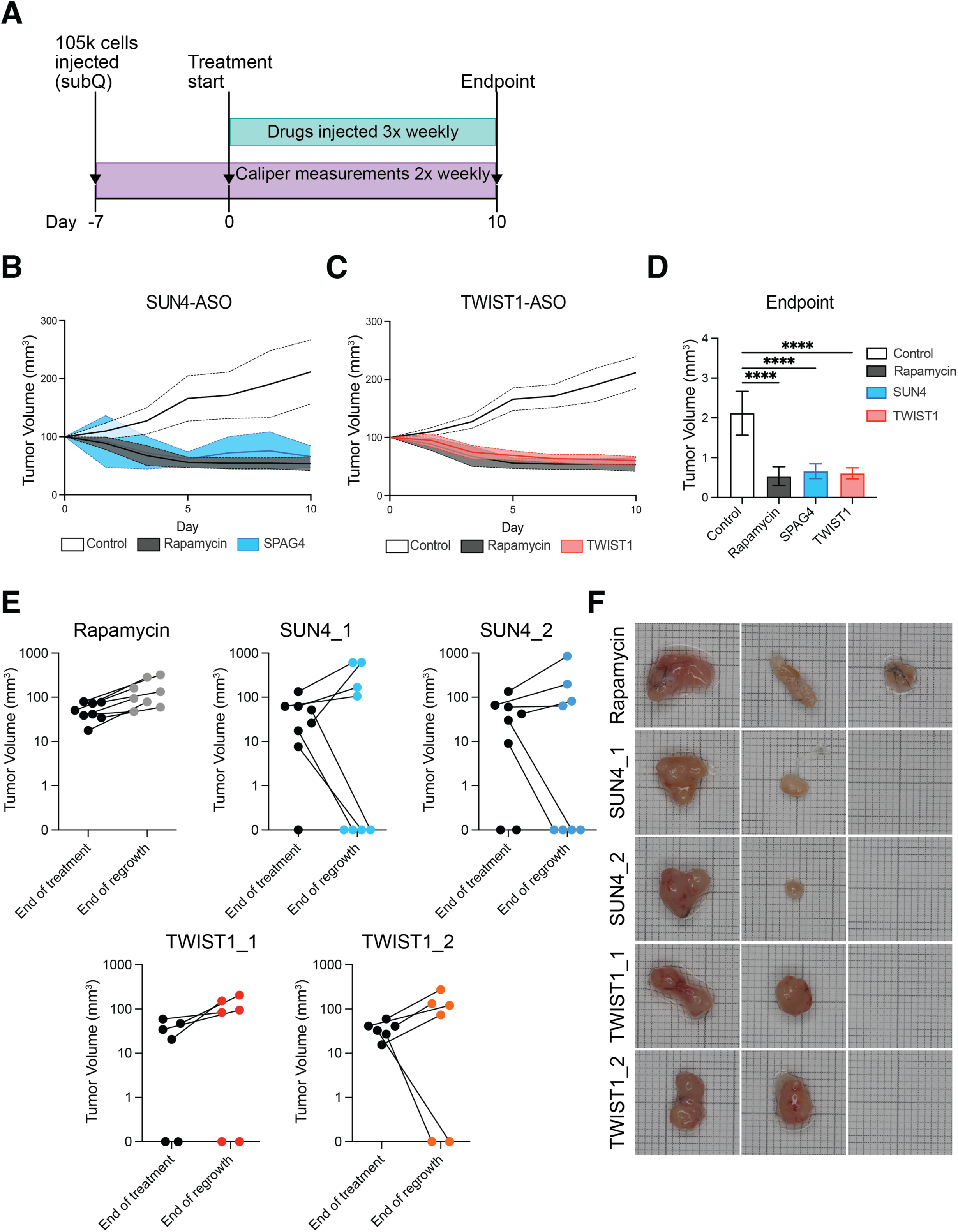
SUN4-ASO and TWIST1-ASO treatment induces complete regression in a subset of tumors *in vivo*. (A) Schematic diagram of treatment strategy. 105k cells were injected subcutaneously, and after the tumor volumes reached 100 mm^3^ (2 tumors per mouse), mice were divided into the following treatment groups: Control (n=4) Rapamycin (1.14 mg/kg, n = 4), SUN4 (1.12mg/kg, n = 4), TWIST1 (1.12mg/kg, n = 4). Animals were treated 3 times a week via i.p. injections. (B&C) Plots of tumor volumes during 10 days of treatment. Solid lines represent mean tumor volume for each group, and shaded regions denote SEM (B: SUN4-ASO, C: TWIST1-ASO). (D) Tumor volume at experimental end point (mean ± SD; n = 4 for each condition; ∗p < 0.05 by using two-way ANOVA and Tukey’s post hoc analysis; Values are normalized to Control). (E) Tumor volume at day 30 (End of treatment) and at experimental endpoint (End of regrowth) for each treatment group. (F) Representative tumors at experimental endpoint for each treatment group.

## Notes

### Competing Interest Statement

The authors have declared no competing interest.

### Summary of Updates

Figures 3-7 and supplemental figures S3-S7 are new while other data has been removed which will be published separately. Due to the major changes, the title has also changed.

## Citations

1. Updated Prevalence of Lymphangioleiomyomatosis in Europe | American Journal of Respiratory and Critical Care Medicine https://www-atsjournals-org.proxy.bib.uottawa.ca/doi/10.1164/rccm.202310-1736LE.

2. Delaney, S.P., Julian, L.M., and Stanford, W.L. (2014). The neural crest lineage as a driver of disease heterogeneity in Tuberous Sclerosis Complex and Lymphangioleiomyomatosis. Front Cell Dev Biol 2. 10.3389/fcell.2014.00069.

3. Salussolia, C.L., Klonowska, K., Kwiatkowski, D.J., and Sahin, M. (2019). Genetic Etiologies, Diagnosis, and Treatment of Tuberous Sclerosis Complex. Annual Review of Genomics and Human Genetics 20, 217–240. 10.1146/annurev-genom-083118-015354.

4. Ryu, J.H., Moss, J., Beck, G.J., Lee, J.-C., Brown, K.K., Chapman, J.T., Finlay, G.A., Olson, E.J., Ruoss, S.J., Maurer, J.R., et al. (2006). The NHLBI Lymphangioleiomyomatosis Registry. Am J Respir Crit Care Med 173, 105–111. 10.1164/rccm.200409-1298OC.

5. De Pauw, R.A., Boelaert, J.R., Haenebalcke, C.W., Matthys, E.G., Schurgers, M.S., and De Vriese, A.S. (2003). Renal angiomyolipoma in association with pulmonary lymphangioleiomyomatosis. American Journal of Kidney Diseases 41, 877–883. 10.1016/S0272-6386(03)00006-4.

6. Bernstein, S.M., Newell, J.D., Jr, Adamczyk, D., Mortenson, R.L., King, T.E., Jr, and Lynch, D.A. (1995). How Common Are Renal Angiomyolipomas in Patients With Pulmonary Lymphangiomyomatosis? Am J Respir Crit Care Med 152, 2138–2143. 10.1164/ajrccm.152.6.8520787.

7. Hayashi, T., Kumasaka, T., Mitani, K., Terao, Y., Watanabe, M., Oide, T., Nakatani, Y., Hebisawa, A., Konno, R., Takahashi, K., et al. (2011). Prevalence of Uterine and Adnexal Involvement in Pulmonary Lymphangioleiomyomatosis: A Clinicopathologic Study of 10 Patients. The American Journal of Surgical Pathology 35, 1776. 10.1097/PAS.0b013e318235edbd.

8. Glasgow, C.G., El-Chemaly, S., and Moss, J. (2012). Lymphatics in lymphangioleiomyomatosis and idiopathic pulmonary fibrosis. European Respiratory Review 21, 196–206. 10.1183/09059180.00009311.

9. Zhe, X., and Schuger, L. (2004). Combined smooth muscle and melanocytic differentiation in lymphangioleiomyomatosis. J Histochem Cytochem 52, 1537–1542. 10.1369/jhc.4A6438.2004.

10. Chu, S.C., Horiba, K., Usuki, J., Avila, N.A., Chen, C.C., Travis, W.D., Ferrans, V.J., and Moss, J. (1999). Comprehensive Evaluation of 35 Patients With Lymphangioleiomyomatosis. Chest 115, 1041–1052. 10.1378/chest.115.4.1041.

11. Müller, N.L., Chiles, C., and Kullnig, P. (1990). Pulmonary lymphangiomyomatosis: correlation of CT with radiographic and functional findings. Radiology 175, 335–339. 10.1148/radiology.175.2.2326457.

12. Lenoir, S., Grenier, P., Brauner, M.W., Frija, J., Remy-Jardin, M., Revel, D., and Cordier, J.F. (1990). Pulmonary lymphangiomyomatosis and tuberous sclerosis: comparison of radiographic and thin-section CT findings. Radiology 175, 329–334. 10.1148/radiology.175.2.2326456.

13. Ryu, J.H., Moss, J., Beck, G.J., Lee, J.-C., Brown, K.K., Chapman, J.T., Finlay, G.A., Olson, E.J., Ruoss, S.J., Maurer, J.R., et al. (2006). The NHLBI Lymphangioleiomyomatosis Registry. Am J Respir Crit Care Med 173, 105–111. 10.1164/rccm.200409-1298OC.

14. Valvezan, A.J., and Manning, B.D. (2019). Molecular logic of mTORC1 signalling as a metabolic rheostat. Nat Metab 1, 321–333. 10.1038/s42255-019-0038-7.

15. Yang, H., Yu, Z., Chen, X., Li, J., Li, N., Cheng, J., Gao, N., Yuan, H.-X., Ye, D., Guan, K.-L., et al. (2021). Structural insights into TSC complex assembly and GAP activity on Rheb. Nat Commun 12, 339. 10.1038/s41467-020-20522-4.

16. Henske, E.P., and McCormack, F.X. (2012). Lymphangioleiomyomatosis — a wolf in sheep’s clothing. J Clin Invest 122, 3807–3816. 10.1172/JCI58709.

17. Bissler, J.J., McCormack, F.X., Young, L.R., Elwing, J.M., Chuck, G., Leonard, J.M., Schmithorst, V.J., Laor, T., Brody, A.S., Bean, J., et al. (2008). Sirolimus for Angiomyolipoma in Tuberous Sclerosis Complex or Lymphangioleiomyomatosis. New England Journal of Medicine 358, 140–151. 10.1056/NEJMoa063564.

18. McCormack, F.X., Inoue, Y., Moss, J., Singer, L.G., Strange, C., Nakata, K., Barker, A.F., Chapman, J.T., Brantly, M.L., Stocks, J.M., et al. (2011). Efficacy and safety of sirolimus in lymphangioleiomyomatosis. N Engl J Med 364, 1595–1606. 10.1056/NEJMoa1100391.

19. Revilla-López, E., Berastegui, C., Méndez, A., Sáez-Giménez, B., Miguel, V.R. de, López-Meseguer, M., Monforte, V., Bravo, C., Pujana, M.A., Ramon, M.A., et al. (2021). Long-term results of sirolimus treatment in lymphangioleiomyomatosis: a single referral centre experience. Scientific Reports 11, 10171. 10.1038/s41598-021-89562-0.

20. Li, J., Kim, S.G., and Blenis, J. (2014). Rapamycin: one drug, many effects. Cell metabolism 19, 373. 10.1016/j.cmet.2014.01.001.

21. Tréfier, A., Tousson-Abouelazm, N., Yamani, L., Ibrahim, S., Joung, K.-B., Pietrobon, A., Yockell-Lelievre, J., Hébert, T.E., Ladak, R.J., Takano, T., et al. (2025). Enhanced Gαq Signaling in TSC2-Deficient Cells Is Required for Their Neoplastic Behavior. Am J Respir Cell Mol Biol 72, 578–590. 10.1165/rcmb.2024-0111OC.

22. Lee, P.-S., Tsang, S.W., Moses, M.A., Trayes-Gibson, Z., Hsiao, L.-L., Jensen, R., Squillace, R., and Kwiatkowski, D.J. (2010). Rapamycin-insensitive up-regulation of MMP2 and other genes in tuberous sclerosis complex 2-deficient lymphangioleiomyomatosis-like cells. Am J Respir Cell Mol Biol 42, 227–234. 10.1165/rcmb.2009-0050OC.

23. Pietrobon, A., Yockell-Lelièvre, J., Melong, N., Smith, L.J., Delaney, S.P., Azzam, N., Xue, C., Merwin, N., Lian, E., Camacho-Magallanes, A., et al. (2023). Tissue-Engineered Disease Modeling of Lymphangioleiomyomatosis Exposes a Therapeutic Vulnerability to HDAC Inhibition. Advanced Science 10, 2302611. 10.1002/advs.202302611.

24. Guo, M., Yu, J.J., Perl, A.K., Wikenheiser-Brokamp, K.A., Riccetti, M., Zhang, E.Y., Sudha, P., Adam, M., Potter, A., Kopras, E.J., et al. (2020). Single-Cell Transcriptomic Analysis Identifies a Unique Pulmonary Lymphangioleiomyomatosis Cell. Am J Respir Crit Care Med 202, 1373–1387. 10.1164/rccm.201912-2445OC.

25. Obraztsova, K., Basil, M.C., Rue, R., Sivakumar, A., Lin, S.M., Mukhitov, A.R., Gritsiuta, A.I., Evans, J.F., Kopp, M., Katzen, J., et al. (2020). mTORC1 activation in lung mesenchyme drives sex- and age-dependent pulmonary structure and function decline. Nat Commun 11, 5640. 10.1038/s41467-020-18979-4.

26. Tang, Y., Kwiatkowski, D.J., and Henske, E.P. (2022). Midkine expression by stem-like tumor cells drives persistence to mTOR inhibition and an immune-suppressive microenvironment. Nat Commun 13, 5018. 10.1038/s41467-022-32673-7.

27. Olatoke, T., Wagner, A., Astrinidis, A., Zhang, E.Y., Guo, M., Zhang, A.G., Mattam, U., Kopras, E.J., Gupta, N., Smith, E.P., et al. (2023). Single-cell multiomic analysis identifies a HOX-PBX gene network regulating the survival of lymphangioleiomyomatosis cells. Science Advances 9, eadf8549. 10.1126/sciadv.adf8549.

28. Goncharova, E.A., Goncharov, D.A., Eszterhas, A., Hunter, D.S., Glassberg, M.K., Yeung, R.S., Walker, C.L., Noonan, D., Kwiatkowski, D.J., Chou, M.M., et al. (2002). Tuberin Regulates p70 S6 Kinase Activation and Ribosomal Protein S6 Phosphorylation: A ROLE FOR THE TSC2 TUMOR SUPPRESSOR GENE INPULMONARY LYMPHANGIOLEIOMYOMATOSIS (LAM)*. Journal of Biological Chemistry 277, 30958–30967. 10.1074/jbc.M202678200.

29. Julian, L.M., Delaney, S.P., Wang, Y., Goldberg, A.A., Doré, C., Yockell-Lelièvre, J., Tam, R.Y., Giannikou, K., McMurray, F., Shoichet, M.S., et al. (2017). Human Pluripotent Stem Cell–Derived TSC2-Haploinsufficient Smooth Muscle Cells Recapitulate Features of Lymphangioleiomyomatosis. Cancer Res 77, 5491–5502. 10.1158/0008-5472.CAN-17-0925.

30. Tam, R.Y., Yockell-Lelièvre, J., Smith, L.J., Julian, L.M., Baker, A.E.G., Choey, C., Hasim, M.S., Dimitroulakos, J., Stanford, W.L., and Shoichet, M.S. (2019). Rationally Designed 3D Hydrogels Model Invasive Lung Diseases Enabling High-Content Drug Screening. Advanced Materials 31, 1806214. 10.1002/adma.201806214.

31. Chambers, S.M., Fasano, C.A., Papapetrou, E.P., Tomishima, M., Sadelain, M., and Studer, L. (2009). Highly efficient neural conversion of human ES and iPS cells by dual inhibition of SMAD signaling. Nat Biotechnol 27, 275–280. 10.1038/nbt.1529.

32. Li, C., Lee, P.-S., Sun, Y., Gu, X., Zhang, E., Guo, Y., Wu, C.-L., Auricchio, N., Priolo, C., Li, J., et al. (2014). Estradiol and mTORC2 cooperate to enhance prostaglandin biosynthesis and tumorigenesis in TSC2-deficient LAM cells. Journal of Experimental Medicine 211, 15–28. 10.1084/jem.20131080.

33. Zhang, H., Hu, Z., Wang, S., Wu, K., Yang, Q., and Song, X. (2022). Clinical features and outcomes of male patients with lymphangioleiomyomatosis: A review. Medicine (Baltimore) 101, e32492. 10.1097/MD.0000000000032492.

34. Muzykewicz, D.A., Sharma, A., Muse, V., Numis, A.L., Rajagopal, J., and Thiele, E.A. (2009). TSC1 and TSC2 mutations in patients with lymphangioleiomyomatosis and tuberous sclerosis complex. Journal of Medical Genetics 46, 465–468. 10.1136/jmg.2008.065342.

35. Martin, K.R., Zhou, W., Bowman, M.J., Shih, J., Au, K.S., Dittenhafer-Reed, K.E., Sisson, K.A., Koeman, J., Weisenberger, D.J., Cottingham, S.L., et al. (2017). The genomic landscape of tuberous sclerosis complex. Nat Commun 8, 15816. 10.1038/ncomms15816.

36. Himes, B.E., Obraztsova, K., Lian, L., Shumyatcher, M., Rue, R., Atochina-Vasserman, E.N., Hur, S.K., Bartolomei, M.S., Evans, J.F., and Krymskaya, V.P. (2018). Rapamycin-independent IGF2 expression in Tsc2-null mouse embryo fibroblasts and human lymphangioleiomyomatosis cells. PLOS ONE 13, e0197105. 10.1371/journal.pone.0197105.

37. Tsc2 disruption in mesenchymal progenitors results in tumors with vascular anomalies overexpressing Lgals3 | eLife https://elifesciences.org/articles/23202.

38. Klover, P.J., Thangapazham, R.L., Kato, J., Wang, J., Anderson, S.A., Hoffmann, V., Steagall, W.K., Li, S., McCart, E., Nathan, N., et al. Tsc2 disruption in mesenchymal progenitors results in tumors with vascular anomalies overexpressing Lgals3. eLife 6, e23202. 10.7554/eLife.23202.

39. Zhao, Z., Rahman, M.A., Chen, Z.G., and Shin, D.M. (2017). Multiple biological functions of Twist1 in various cancers. Oncotarget 8, 20380–20393. 10.18632/oncotarget.14608.

40. Feng, W., Wang, G., Zhang, X., Cheng, S., Song, X., Zhang, P., and Wang, J. (2022). New understanding of SPAG4 as a tumor marker: SPAG4 is a potential pan-cancer biomarker for prognosis and immunotherapy. Preprint at In Review, 10.21203/rs.3.rs-1704715/v1 https://doi.org/10.21203/rs.3.rs-1704715/v1.

41. Wang, Y., Tang, Y., Li, J., Wang, D., and Li, W. (2021). Human sperm-associated antigen 4 as a potential prognostic biomarker of lung squamous cell carcinoma. J Int Med Res 49, 03000605211032807. 10.1177/03000605211032807.

42. Pietrobon, A., Yockell-Lelièvre, J., Flood, T.A., and Stanford, W.L. (2022). Renal organoid modeling of tuberous sclerosis complex reveals lesion features arise from diverse developmental processes. Cell Reports 40, 111048. 10.1016/j.celrep.2022.111048.

43. Ji, Y., Jiang, J., Huang, L., Feng, W., Zhang, Z., Jin, L., and Xing, X. (2018). Sperm-associated antigen 4 (SPAG4) as a new cancer marker interacts with Nesprin3 to regulate cell migration in lung carcinoma. Oncol Rep. 10.3892/or.2018.6473.

44. Thoma, H., Grünewald, L., Braune, S., Pasch, E., and Alsheimer, M. (2023). SUN4 is a spermatid type II inner nuclear membrane protein that forms heteromeric assemblies with SUN3 and interacts with lamin B3. J Cell Sci 136, jcs260155. 10.1242/jcs.260155.

45. Fan, X., Masamsetti, V.P., Sun, J.Q., Engholm-Keller, K., Osteil, P., Studdert, J., Graham, M.E., Fossat, N., and Tam, P.P. (2021). TWIST1 and chromatin regulatory proteins interact to guide neural crest cell differentiation. eLife. 10.7554/eLife.62873.

46. Zheng, Y., Dai, M., Dong, Y., Yu, H., Liu, T., Feng, X., Yu, B., Zhang, H., Wu, J., Kong, W., et al. (2022). ZEB2/TWIST1/PRMT5/NuRD Multicomplex Contributes to the Epigenetic Regulation of EMT and Metastasis in Colorectal Carcinoma. Cancers 14, 3426. 10.3390/cancers14143426.

47. Ren, J., and Crowley, S.D. (2020). Twist1: A Double-Edged Sword in Kidney Diseases. KDD 6, 247–257. 10.1159/000505188.

48. Parkhitko, A.A., Priolo, C., Coloff, J.L., Yun, J., Wu, J.J., Mizumura, K., Xu, W., Malinowska, I.A., Yu, J., Kwiatkowski, D.J., et al. (2014). Autophagy-Dependent Metabolic Reprogramming Sensitizes TSC2-Deficient Cells to the Antimetabolite 6-Aminonicotinamide. Mol Cancer Res 12, 48–57. 10.1158/1541-7786.MCR-13-0258-T.

49. Collotta, D., Bertocchi, I., Chiapello, E., and Collino, M. (2023). Antisense oligonucleotides: a novel Frontier in pharmacological strategy. Front. Pharmacol. 14, 1304342. 10.3389/fphar.2023.1304342.

50. Lightfoot, H., Schneider, A., and Hall, J. (2018). Pharmacokinetics and Pharmacodynamics of Antisense Oligonucleotides. In Oligonucleotide-Based Drugs and Therapeutics, N. Ferrari and R. Seguin, eds. (Wiley), pp. 107–136. 10.1002/9781119070153.ch4.

51. Bissler, J.J., McCormack, F.X., Young, L.R., Elwing, J.M., Chuck, G., Leonard, J.M., Schmithorst, V.J., Laor, T., Brody, A.S., Bean, J., et al. (2008). Sirolimus for Angiomyolipoma in Tuberous Sclerosis Complex or Lymphangioleiomyomatosis. New England Journal of Medicine 358, 140–151. 10.1056/NEJMoa063564.

52. Renal manifestation of tuberous sclerosis complex - Bissler - 2018 - American Journal of Medical Genetics Part C: Seminars in Medical Genetics - Wiley Online Library https://onlinelibrary-wiley-com.proxy.bib.uottawa.ca/doi/10.1002/ajmg.c.31654.

53. Yue, M., Pacheco, G., Cheng, T., Li, J., Wang, Y., Henske, E.P., and Schuger, L. (2016). Evidence Supporting a Lymphatic Endothelium Origin for Angiomyolipoma, a TSC2− Tumor Related to Lymphangioleiomyomatosis. Am J Pathol 186, 1825–1836. 10.1016/j.ajpath.2016.03.009.

54. Obraztsova, K., Basil, M.C., Rue, R., Sivakumar, A., Lin, S.M., Mukhitov, A.R., Gritsiuta, A.I., Evans, J.F., Kopp, M., Katzen, J., et al. (2020). mTORC1 activation in lung mesenchyme drives sex- and age-dependent pulmonary structure and function decline. Nat Commun 11, 5640. 10.1038/s41467-020-18979-4.

55. Pietrobon, A., Yockell-Lelièvre, J., Flood, T.A., and Stanford, W.L. (2022). Renal organoid modeling of tuberous sclerosis complex reveals lesion features arise from diverse developmental processes. Cell Reports 40, 111048. 10.1016/j.celrep.2022.111048.

56. Stundl, J., Desingu Rajan, A.R., and Bronner, M.E. (2026). Neural crest gene regulatory networks as drivers of development, diversification and disease. Nat Rev Mol Cell Biol, 1–19. 10.1038/s41580-026-00949-1.

57. Brunelli, A., Catalini, G., and Fianchini, A. (1996). Pregnancy exacerbating unsuspected mediastinal lymphangioleiomyomatosis and chylothorax. Int J Gynaecol Obstet 52, 289–290.

58. Yano, S. (2002). Exacerbation of pulmonary lymphangioleiomyomatosis by exogenous oestrogen used for infertility treatment. Thorax 57, 1085–1086.

59. Shen, A., Iseman, M.D., Waldron, J.A., and King, T.E. (1987). Exacerbation of Pulmonary Lymphangioleiomyomatosis by Exogenous Estrogens. Chest 91, 782–785. 10.1378/chest.91.5.782.

60. Oberstein, E.M., Fleming, L.E., Gómez-Marin, O., and Glassberg, M.K. (2003). Pulmonary Lymphangioleiomyomatosis (LAM): Examining Oral Contraceptive Pills and the Onset of Disease. Journal of Women’s Health 12, 81–85. 10.1089/154099903321154176.

61. Gupta, N., Lee, H.-S., Ryu, J.H., Taveira-DaSilva, A.M., Beck, G.J., Lee, J.-C., McCarthy, K., Finlay, G.A., Brown, K.K., Ruoss, S.J., et al. (2018). The NHLBI LAM Registry: Prognostic Physiologic and Radiologic Biomarkers Emerge From a 15-Year Prospective Longitudinal Analysis. Chest. 10.1016/j.chest.2018.06.016.

62. Berger, U., Khaghani, A., Pomerance, A., Yacoub, M.H., and Coombes, R.C. (1990). Pulmonary lymphangioleiomyomatosis and steroid receptors. An immunocytochemical study. Am. J. Clin. Pathol. 93, 609–614.

63. Ohori, N.P., Yousem, S.A., Sonmez-Alpan, E., and Colby, T.V. (1991). Estrogen and progesterone receptors in lymphangioleiomyomatosis, epithelioid hemangioendothelioma, and sclerosing hemangioma of the lung. Am. J. Clin. Pathol. 96, 529–535.

64. Matsui, K., Takeda, K., Yu, Z.X., Valencia, J., Travis, W.D., Moss, J., and Ferrans, V.J. (2000). Downregulation of estrogen and progesterone receptors in the abnormal smooth muscle cells in pulmonary lymphangioleiomyomatosis following therapy. An immunohistochemical study. Am. J. Respir. Crit. Care Med. 161, 1002–1009. 10.1164/ajrccm.161.3.9904009.

65. Prizant, H., Sen, A., Light, A., Cho, S.-N., DeMayo, F.J., Lydon, J.P., and Hammes, S.R. (2013). Uterine-specific loss of Tsc2 leads to myometrial tumors in both the uterus and lungs. Mol. Endocrinol. 27, 1403–1414. 10.1210/me.2013-1059.

66. Yu, J., Astrinidis, A., Howard, S., and Henske, E.P. (2004). Estradiol and tamoxifen stimulate LAM-associated angiomyolipoma cell growth and activate both genomic and nongenomic signaling pathways. Am. J. Physiol. Lung Cell Mol. Physiol. 286, L694–700. 10.1152/ajplung.00204.2003.

67. Yu, J.J., Robb, V.A., Morrison, T.A., Ariazi, E.A., Karbowniczek, M., Astrinidis, A., Wang, C., Hernandez-Cuebas, L., Seeholzer, L.F., Nicolas, E., et al. (2009). Estrogen promotes the survival and pulmonary metastasis of tuberin-null cells. Proc. Natl. Acad. Sci. U.S.A. 106, 2635–2640. 10.1073/pnas.0810790106.

68. Gu, X., Yu, J.J., Ilter, D., Blenis, N., Henske, E.P., and Blenis, J. (2013). Integration of mTOR and estrogen–ERK2 signaling in lymphangioleiomyomatosis pathogenesis. PNAS, 201309110. 10.1073/pnas.1309110110.

69. Yano, S. (2002). Exacerbation of pulmonary lymphangioleiomyomatosis by exogenous oestrogen used for infertility treatment. Thorax 57, 1085–1086. 10.1136/thorax.57.12.1085.

70. Taveira-DaSilva, A.M., Stylianou, M.P., Hedin, C.J., Hathaway, O., and Moss, J. (2004). Decline in lung function in patients with lymphangioleiomyomatosis treated with or without progesterone. Chest 126, 1867–1874. 10.1378/chest.126.6.1867.

71. Tai, J., Liu, S., Yan, X., Huang, L., Pan, Y., Huang, H., Zhao, Z., Xu, B., and Liu, J. (2024). Novel developments in the study of estrogen in the pathogenesis and therapeutic intervention of lymphangioleiomyomatosis. Orphanet J Rare Dis 19, 236. 10.1186/s13023-024-03239-1.

72. Taveira-DaSilva, A.M., Hathaway, O., Stylianou, M., and Moss, J. (2011). Changes in Lung Function and Chylous Effusions in Patients With Lymphangioleiomyomatosis Treated With Sirolimus. Ann Intern Med 154, 797-292–293. 10.1059/0003-4819-154-12-201106210-00007.

73. Dabora, S.L., Franz, D.N., Ashwal, S., Sagalowsky, A., DiMario, F.J., Miles, D., Cutler, D., Krueger, D., Uppot, R.N., Rabenou, R., et al. (2011). Multicenter Phase 2 Trial of Sirolimus for Tuberous Sclerosis: Kidney Angiomyolipomas and Other Tumors Regress and VEGF- D Levels Decrease. PLoS One 6, e23379. 10.1371/journal.pone.0023379.

74. Bee, J., Fuller, S., Miller, S., and Johnson, S.R. (2018). Lung function response and side effects to rapamycin for lymphangioleiomyomatosis: a prospective national cohort study. Thorax 73, 369–375. 10.1136/thoraxjnl-2017-210872.

75. McCormack, F.X., Inoue, Y., Moss, J., Singer, L.G., Strange, C., Nakata, K., Barker, A.F., Chapman, J.T., Brantly, M.L., Stocks, J.M., et al. (2011). Efficacy and Safety of Sirolimus in Lymphangioleiomyomatosis. N Engl J Med 364, 1595–1606. 10.1056/NEJMoa1100391.

76. Revilla-López, E., Berastegui, C., Méndez, A., Sáez-Giménez, B., Ruiz de Miguel, V., López-Meseguer, M., Monforte, V., Bravo, C., Pujana, M.A., Ramon, M.A., et al. (2021). Long-term results of sirolimus treatment in lymphangioleiomyomatosis: a single referral centre experience. Sci Rep 11, 10171. 10.1038/s41598-021-89562-0.

77. Qin, Q., Xu, Y., He, T., Qin, C., and Xu, J. (2012). Normal and disease-related biological functions of Twist1 and underlying molecular mechanisms. Cell Res 22, 90– 106. 10.1038/cr.2011.144.

78. Zhao, Z., Rahman, M.A., Chen, Z.G., and Shin, D.M. (2017). Multiple biological functions of Twist1 in various cancers. Oncotarget 8, 20380–20393. 10.18632/oncotarget.14608.

79. Jin, H.-O., Hong, S.-E., Woo, S.-H., Lee, J.-H., Choe, T.-B., Kim, E.-K., Noh, W.-C., Lee, J.-K., Hong, S.-I., Kim, J.-I., et al. (2012). Silencing of Twist1 sensitizes NSCLC cells to cisplatin via AMPK-activated mTOR inhibition. Cell Death Dis 3, e319. 10.1038/cddis.2012.63.

80. Lv, T., Wang, Q., Cromie, M., Liu, H., Tang, S., Song, Y., and Gao, W. (2015). Twist1-mediated 4E-BP1 regulation through mTOR in non-small cell lung cancer. Oncotarget 6, 33006–33018.

81. Marcucci, F., Stassi, G., and De Maria, R. (2016). Epithelial–mesenchymal transition: a new target in anticancer drug discovery. Nat Rev Drug Discov 15, 311–325. 10.1038/nrd.2015.13.

82. McGillivary, R.M., Starr, D.A., and Luxton, G.W.G. (2023). Building and breaking mechanical bridges between the nucleus and cytoskeleton: Regulation of LINC complex assembly and disassembly. Current Opinion in Cell Biology 85, 102260. 10.1016/j.ceb.2023.102260.

83. Hornberger, T.A. (2011). Mechanotransduction and the Regulation of mTORC1 Signaling in Skeletal Muscle. Int J Biochem Cell Biol 43, 1267–1276. 10.1016/j.biocel.2011.05.007.

84. Di, X., Gao, X., Peng, L., Ai, J., Jin, X., Qi, S., Li, H., Wang, K., and Luo, D. (2023). Cellular mechanotransduction in health and diseases: from molecular mechanism to therapeutic targets. Sig Transduct Target Ther 8, 282. 10.1038/s41392-023-01501-9.

85. Maurer, M., and Lammerding, J. (2019). The Driving Force: Nuclear Mechanotransduction in Cellular Function, Fate, and Disease. Annu Rev Biomed Eng 21, 443–468. 10.1146/annurev-bioeng-060418-052139.

86. Zhu, L., Muhtar, P., Goyama, S., and Yoshimi, A. (2026). Advances in antisense oligonucleotide treatment for cancer. Jpn J Clin Oncol, hyag017. 10.1093/jjco/hyag017.

87. Chen, J., Nie, L., Lee, S., Chu, H., Tian, H., Wang, Y., He, W., Jemielita, T., Gruber, S., Song, Y., et al. (2025). Challenges and Possible Strategies to Address Them in Rare Disease Drug Development: A Statistical Perspective. Clinical Pharmacology & Therapeutics 118, 62–73. 10.1002/cpt.3631.

88. Ziegler, A., Carroll, J., Bain, J.M., Sands, T.T., Fee, R.J., Uher, D., Kanner, C.H., Montes, J., Glass, S., Douville, J., et al. (2024). Antisense oligonucleotide therapy in an individual with KIF1A-associated neurological disorder. Nat Med 30, 2782–2786. 10.1038/s41591-024-03197-y.

89. Cheerie, D., Meserve, M.M., Beijer, D., Kaiwar, C., Newton, L., Tavares, A.L.T., Verran, A.S., Sherrill, E., Leonard, S., Sanders, S.J., et al. (2025). Consensus guidelines for assessing eligibility of pathogenic DNA variants for antisense oligonucleotide treatments. The American Journal of Human Genetics 112, 975–983. 10.1016/j.ajhg.2025.02.017.

90. Kim, J., Woo, S., de Gusmao, C.M., Zhao, B., Chin, D.H., DiDonato, R.L., Nguyen, M.A., Nakayama, T., Hu, C.A., Soucy, A., et al. (2023). A framework for individualized splice-switching oligonucleotide therapy. Nature 619, 828–836. 10.1038/s41586-023-06277-0.

91. Kim, J., Hu, C., Moufawad El Achkar, C., Black, L.E., Douville, J., Larson, A., Pendergast, M.K., Goldkind, S.F., Lee, E.A., Kuniholm, A., et al. (2019). Patient-Customized Oligonucleotide Therapy for a Rare Genetic Disease. N Engl J Med 381, 1644–1652. 10.1056/NEJMoa1813279.

92. Ersöz, E., and Demir-Dora, D. (2024). Unveiling the potential of antisense oligonucleotides: Mechanisms, therapies, and safety insights. Drug Development Research 85, e22187. 10.1002/ddr.22187.

93. Talukdar, P.D., and Chatterji, U. (2023). Transcriptional co-activators: emerging roles in signaling pathways and potential therapeutic targets for diseases. Sig Transduct Target Ther 8, 427. 10.1038/s41392-023-01651-w.

94. Su, B.G., and Henley, M.J. (2021). Drugging Fuzzy Complexes in Transcription. Front Mol Biosci 8, 795743. 10.3389/fmolb.2021.795743.

95. Méjat, A., and Misteli, T. (2010). LINC complexes in health and disease. Nucleus 1, 40–52. 10.4161/nucl.1.1.10530.

96. Chiarini, F., Paganelli, F., Balestra, T., Capanni, C., Fazio, A., Manara, M.C., Landuzzi, L., Petrini, S., Evangelisti, C., Lollini, P.-L., et al. (2022). Lamin A and the LINC complex act as potential tumor suppressors in Ewing Sarcoma. Cell Death Dis 13, 346. 10.1038/s41419-022-04729-5.

97. Hansen, E., and Holaska, J.M. (2023). The nuclear envelope and metastasis. Oncotarget 14, 317–320. 10.18632/oncotarget.28375.

98. Cheng, Q., Wei, T., Farbiak, L., Johnson, L.T., Dilliard, S.A., and Siegwart, D.J. (2020). Selective ORgan Targeting (SORT) nanoparticles for tissue specific mRNA delivery and CRISPR/Cas gene editing. Nat Nanotechnol 15, 313–320. 10.1038/s41565-020-0669-6.

99. Navid Talemi, M., Ramezani Farani, M., Alipour Eskandani, N., Mirzaee, D., Alipourfard, I., and Huh, Y.S. (2026). Programmable lipid nanoparticles for RNA therapeutics: Design principles and clinical translation. Materials Today Bio 37, 102774. 10.1016/j.mtbio.2026.102774.

100. Teratoma Formation: A Tool for Monitoring Pluripotency in Stem Cell Research - PMC https://pmc-ncbi-nlm-nih-gov.proxy.bib.uottawa.ca/articles/PMC4402211/.

101. Mehlem, A., Hagberg, C.E., Muhl, L., Eriksson, U., and Falkevall, A. (2013). Imaging of neutral lipids by oil red O for analyzing the metabolic status in health and disease. Nat Protoc 8, 1149–1154. 10.1038/nprot.2013.055.

102. Kolberg, L., Raudvere, U., Kuzmin, I., Adler, P., Vilo, J., and Peterson, H. (2023). g:Profiler—interoperable web service for functional enrichment analysis and gene identifier mapping (2023 update). Nucleic Acids Research 51, W207–W212. 10.1093/nar/gkad347.

103. Sanjana, N.E., Shalem, O., and Zhang, F. (2014). Improved vectors and genome-wide libraries for CRISPR screening. Nat Methods 11, 783–784. 10.1038/nmeth.3047.

104. Sanjana, N.E., Shalem, O., and Zhang, F. (2014). Improved vectors and genome-wide libraries for CRISPR screening. Nat Methods 11, 783–784. 10.1038/nmeth.3047.

105. Hart, T., and Moffat, J. (2016). BAGEL: a computational framework for identifying essential genes from pooled library screens. BMC Bioinformatics 17, 164. 10.1186/s12859-016-1015-8.

106. Shalem, O., Sanjana, N.E., Hartenian, E., Shi, X., Scott, D.A., Mikkelsen, T.S., Heckl, D., Ebert, B.L., Root, D.E., Doench, J.G., et al. (2014). Genome-Scale CRISPR-Cas9 Knockout Screening in Human Cells. Science 343, 84–87. 10.1126/science.1247005.

107. Li, W., Köster, J., Xu, H., Chen, C.-H., Xiao, T., Liu, J.S., Brown, M., and Liu, X.S. (2015). Quality control, modeling, and visualization of CRISPR screens with MAGeCK-VISPR. Genome Biology 16, 281. 10.1186/s13059-015-0843-6.

108. Wang, B., Wang, M., Zhang, W., Xiao, T., Chen, C.-H., Wu, A., Wu, F., Traugh, N., Wang, X., Li, Z., et al. (2019). Integrative analysis of pooled CRISPR genetic screens using MAGeCKFlute. Nat Protoc 14, 756–780. 10.1038/s41596-018-0113-7.

109. Maritan, S.M., Lian, E.Y., and Mulligan, L.M. (2017). An Efficient and Flexible Cell Aggregation Method for 3D Spheroid Production. JoVE, 55544. 10.3791/55544.

